# A novel computational complete deconvolution method using RNA-seq data

**DOI:** 10.1101/496596

**Authors:** Kai Kang, Qian Meng, Igor Shats, David M. Umbach, Melissa Li, Yuanyuan Li, Xiaoling Li, Leping Li

**Affiliations:** Biostatistics and Computational Biology Branch, NIEHS/NIH; Signal Transduction Laboratory, NIEHS/NIH

## Abstract

The cell type composition of many biological tissues varies widely across samples. Such sample heterogeneity hampers efforts to probe the role of each cell type in the tissue microenvironment. Current approaches that address this issue have drawbacks. Cell sorting or single-cell based experimental techniques disrupt in situ interactions and alter physiological status of cells in tissues. Computational methods are flexible and promising; but they often estimate either sample-specific proportions of each cell type or cell-type-specific gene expression profiles, not both, by requiring the other as input. We introduce a computational **C**omplete **D**econvolution method that can estimate both sample-specific proportions of each cell type and cell-type-specific gene expression profiles simultaneously using bulk RNA-**Seq** data only (CDSeq). We assessed our method’s performance using several synthetic and experimental mixtures of varied but known cell type composition and compared its performance to the performance of two state-of-the art deconvolution methods on the same mixtures. The results showed CDSeq can estimate both sample-specific proportions of each component cell type and cell-typespecificgene expression profiles with high accuracy. CDSeq holds promise for computationally deciphering complex mixtures of cell types, each with differing expression profiles, using RNA-seq data measured in bulk tissue (MATLAB code is available at https://github.com/kkang7/CDSeq_011).

## Introduction

Researchers measure gene expression in bulk tissue samples to gain insight into how biological processes change through development or under differing experimental conditions. A tissue, such as a liver or a tumor, is, however, heterogeneous, comprising multiple cell types that vary in their proportional contribution from sample to sample. The measured expression of a gene in a bulk sample reflects the expression of that gene in each cell in the sample. Consequently, the measured gene expression profile (GEP) of a tissue sample is commonly regarded as a weighted average of the GEPs of the different component cell types^1,2^.

The heterogenous nature of bulk tissue samples complicates the interpretation of bulk measurements such as RNA-seq. Often researchers are interested in understanding whether an experimental treatment targets one particular cell type in a heterogeneous tissue or to investigate possible sources of variation among samples^3^. For example, the composition of tumor-infiltrating lymphocytes impacts the tumor growth and patients’ clinical outcomes^4–9^. To those ends, understanding the cell-type composition of each sample and the GEP of each constituent cell type becomes important. “Deconvolution” is a generic term for a procedure that estimates the proportion of each cell type in a bulk sample together with their corresponding cell-type-specific GEPs^10,11^. Deconvolution can be approached experimentally using flow cytometry or single cell RNA sequencing. For solid tissues, these techniques require separating individual cells, thereby presenting laboratory challenges as well as potentially sacrificing a systems perspective. Single cell RNA sequencing is also expensive and requires challenging data handling and analysis ^12,13^.

Deconvolution can also be approached computationally using GEP profiles from collections of bulk tissue samples^10,14^. With a wealth of available next generation sequencing (NGS) gene expression data, computational methods of deconvolution become increasingly desirable. Several deconvolution methods have been developed in the past decade. The pioneering work of Venet et al.^15^ employed an algorithm based on matrix factorization to deconvolve a matrix of GEPs (each normalized to sum to 1) into a product of two matrices, one containing the cell-type proportions for each sample and the other containing the GEPs for each cell type. The constraints required for each matrix in the product (proportions must be non-negative and sum to 1 across cell types; expression levels must obey the same constraints across genes) impose technical challenges on matrix factorization in this context. Consequently, most existing methods only perform partial deconvolution: either the algorithms require cell-type proportions as input to estimate cell-type-specific GEPs^1,16–18^ or vice versa^19–23^. These methods generally use regression techniques and some also use marker genes^3,24,25^ to estimate the unknowns of interest. Such partial deconvolution approaches have shown important findings^9,24^, however they can suffer if the needed information is unavailable or if the fidelity of reference profiles or proportions is questionable. In addition, most deconvolution algorithms report cell-type proportions based on the proportion of RNA that each cell type contributes to total RNA from the bulk sample; whereas researchers are often more interested in the proportion of cells of each type in the bulk sample. These proportions are equivalent only when all cell types contribute about the same amount of RNA per cell.

We developed a complete deconvolution algorithm for bulk RNA-seq data from heterogeneous tissue samples. Our goal was to improve on partial deconvolution by estimating cell-type proportions and cell-type-specific GEPs simultaneously and in an unsupervised fashion. Our algorithm was inspired by latent Dirichlet allocation (LDA)^26^, a probabilistic model designed for natural language processing, that also inspired several existing partial deconvolution methods ^19,20,27^. The original LDA model does not, however, fully capture the complexity of bulk RNA-seq data. Our proposed model extended the original LDA model in two primary ways that would be unnecessary in the context of natural language processing but are crucial for RNA-seq data. First, we built in a dependence of gene expression on gene length. Second, we accommodated possibly different amounts of RNA per cell from cell types whose cells differ in size when estimating the proportion of cells of each type in the sample^28^. The cell types constructed by our algorithm are anonymous and must be identified via comparison to reference cell-type-specific GEPs. In addition, instead of specifying the number of cell types a priori, we provide an algorithm that allows the data to guide selection of the number of cell types.

Finally, we proposed a quasi-unsupervised learning strategy that augments the input data (GEPs from mixed samples) with additional GEPs from pure cell lines that are anticipated to be components of the mixture. We examined whether this strategy can improve the algorithm’s performance in some complex situations where a fully unsupervised learning strategy yields difficult-to-interpret solutions.

Ideally, we would compare the performance of our complete deconvolution method (CDSeq) to other complete deconvolution methods; however, few^15,29^ have been published. Among those, either links to the code provided are outdated^15^ or the method requires generally unavailable information about marker genes for constituent cell types^29^. Therefore, we tested the performance of CDSeq in comparison to two partial deconvolution methods: CIBERSORT^23^, which estimates cell-type proportions based on relative amounts for RNA when cell-type-specific GEPs are provided; and csSAM^1^, which estimates cell-type-specific GEPs when sample-specific cell-type proportions are provided. These comparisons encompassed a range of data sets: synthetic mixtures created numerically from GEPs of pure cell lines downloaded from the Cold Spring Harbor Laboratory (CHSL) website, GEPs measured on heterogenous RNA samples constructed in our lab by mixing RNA extracted from pure cell lines in different proportions, the experimental expression data that was used to evaluate csSAM^1^, expression data from follicular lymphoma samples^23^, and expression data from samples of peripheral blood mononuclear cells (PBMC)^23^. In all these comparisons, CDSeq performed as well or better than competitors in providing accurate estimates of cell-type proportions and cell-type-specific GEPs from heterogeneous tissue samples.

## Results

CDSeq is a complete deconvolution algorithm that takes RNA-seq data (raw read counts) from a collection of possibly heterogeneous samples as input and returns estimates of GEPs for each constituent cell type as well as the proportional representation of those cell types (schematic in **Figure 1**; details of algorithm in Methods). We evaluated our estimation of cell-type proportions in comparison to CIBERSORT^23^, which requires cell-type-specific GEPs as input. When GEPs of pure cell lines that constitute the heterogeneous samples were available, we provided them to CIBERSORT. We evaluated our estimation of cell-type-specific GEPs in comparison to csSAM^1^, which requires cell-type proportions as input. The proportion information provided to csSAM was either known in advance or estimated using flow cytometry. To mitigate the differences of required inputs between CIBERSORT, csSAM, which are mostly applied for microarray data, and CDSeq, which is designed for RNA-seq data, we provided the RPKM normalization instead of raw read counts data to CIBERSORT and csSAM.

**Figure 1.**
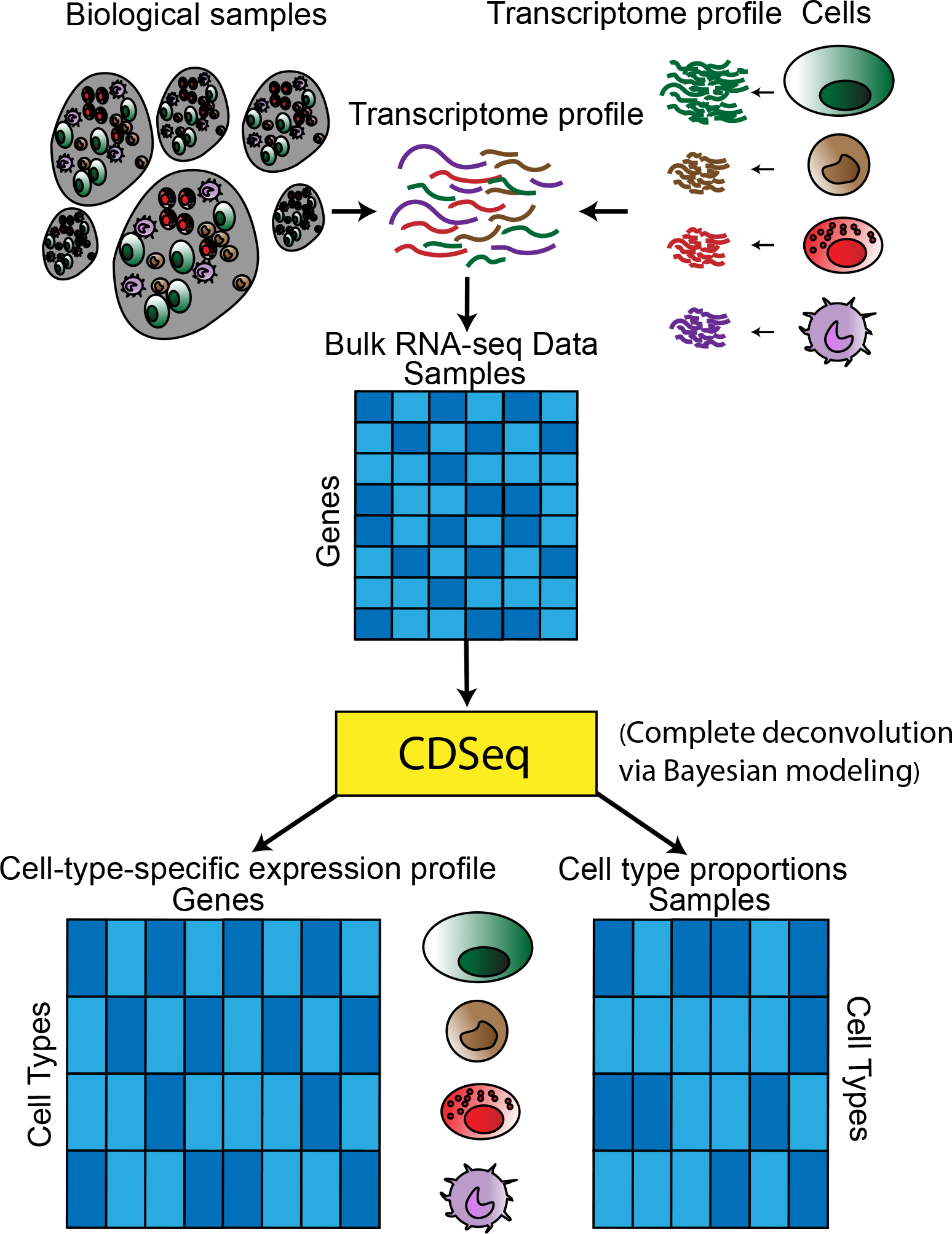
Schematic of the CDSeq approach. Heterogeneous samples consist of different cell types. The cell size of each cell type may differ across types, as the green cell depicted is bigger and consequently produces more RNA than the other cell types. The bulk RNA-seq profile from biological samples represents a weighted average of the expression profiles of the constituent cell types. CDSeq takes as input the bulk RNA-seq data for a collection of samples and performs complete deconvolution that outputs estimates of both the cell-type-specific expression profiles and the cell-type proportions for each sample. To identify the estimated cell types, one can compare the estimated GEPs to reference GEPs (Methods). This figure depicts a simple scenario of six biological samples comprising four cell types, each with gene expression measurements on eight genes.

### Performance on synthetic data

We first benchmarked CDSeq on synthetic mixtures with known composition that we created numerically from publicly available GEPs from CSHL (Methods). In this synthetic numerical experiment, we amplified the potential bias between RNA proportions and cell-type proportions by artificially increasing the RNA amount of certain cell types before mixing them together to generate the synthetic samples. We generated 40 synthetic samples containing six cell types mixed in differing proportions (Methods).

In estimating cell-type proportions, CDSeq outperformed CIBERSORT, showing smaller differences between the true and estimated proportions for each cell type and, consequently, smaller root mean square error (RMSE) (Methods) (**Figure 2** and **Supplementary Figure 1**). The RMSE of CDSeq was 77% lower than that of CIBERSORT, particularly for normal mammary epithelial cells, a cell type where our synthetic cells contributed twice the RNA as cells of most other types so that estimates of the proportion of cells as provided by CDSeq were expected to deviate from estimates of proportion of RNA as provided by CIBERSORT.

**Figure 2.**
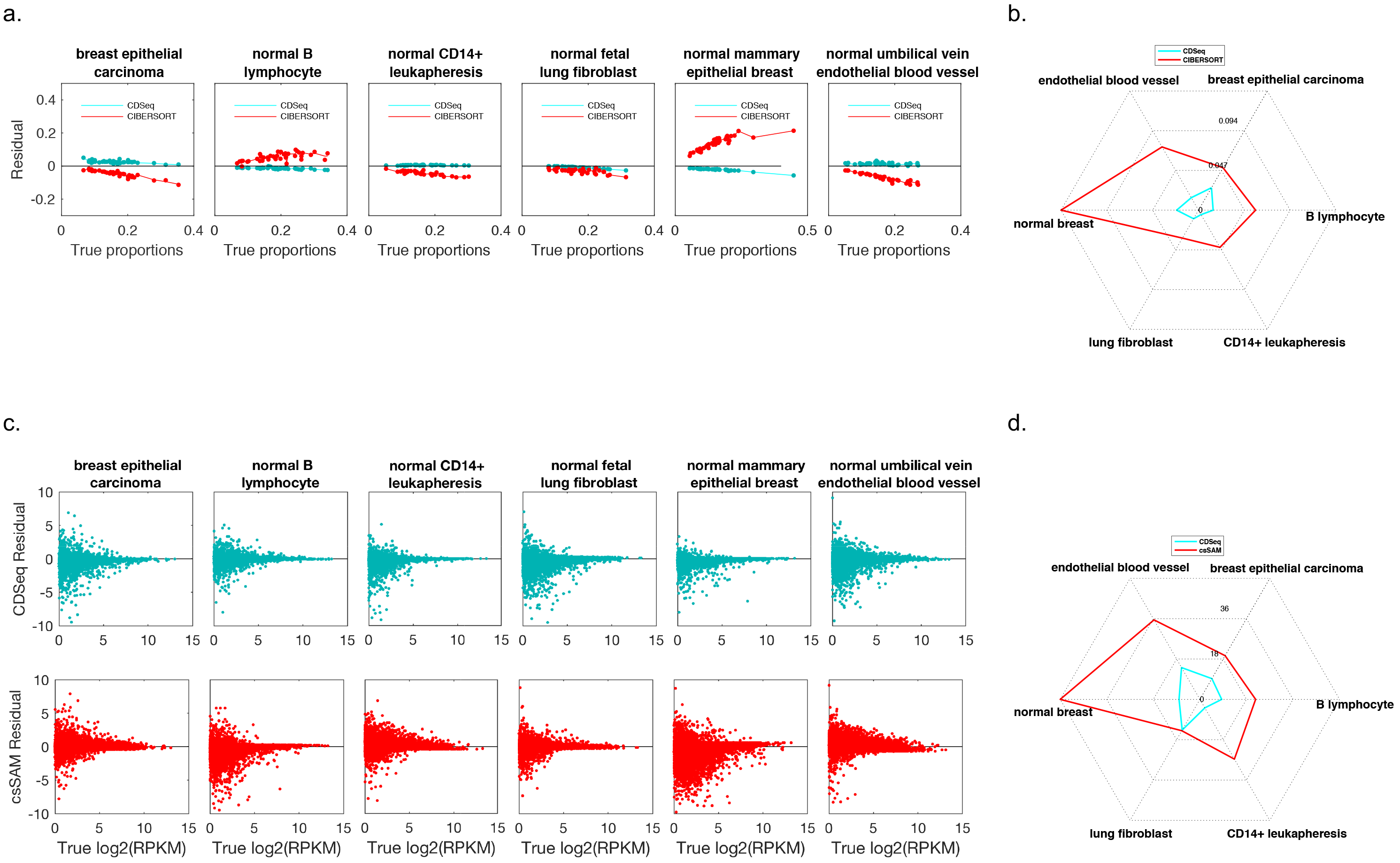
Deconvolution of synthetic mixtures. We used expression data for six pure cell lines downloaded from CSHL to produce data from 40 synthetic mixtures with known cell-type proportions for testing the performance of CDSeq. (**Supplementary Table 1**). To artificially amplify the confounding factor of cell size differences, we multiplied the cell line read counts by a predefined vector to double RNA-seq read counts in the normal B lymphocyte and normal mammary epithelial breast components (chosen at random). We ran: CDSeq with six cell types, *α* = 5, *β* = 0.5 and 700 MCMC runs; CIBERSORT with the default settings to estimate proportions while providing pure cell line expression profiles as input; and csSAM with the default settings to estimate cell-type-specific GEPs while providing true sample-specific cell-type proportions as input. (a) Difference (“residual”) between estimated and true cell-type proportion plotted against true proportion for CDSeq (green) and CIBERSORT (red). Each plotted point represents the value for a single sample. Lowess smooth was added to aid comparison. (b) Radar plot of RMSE for estimates of sample-specific cell-type proportions. CDSeq (green); CIBERSORT (red). Total RMSE summing over cell types is 77% smaller for CDSeq compared to CIBERSORT. (c) Difference (“residual”) between estimated and true log_2_ gene expression level [(log_2_(RPKM)] plotted against true log_2_ gene expression level for CDseq (green) and csSAM (red). Each plotted point represents a single gene, 22498 genes total. (d) Radar plot of RMSE for gene expression levels (RPKM). CDSeq (green); csSAM (red). Total RMSE of gene expression (summing over cell types) is 64% smaller for CDSeq compared to csSAM.

In estimating GEPs, CDSeq outperformed csSAM, though performance of each was comparable. At low expression levels (log_2_ (RPKM)<5) neither method provided estimates that were accurate and precise; however, at higher expression levels, CDSeq estimates tended to be somewhat closer to true expression level than csSAM estimates, resulting in lower RMSE values for CDSeq estimates of expression in every cell type (**Figure 2** and **Supplementary Figure 1**).

### Performance on RNA extracted from cultured cells and mixed

Our second performance evaluation used data from a designed experiment that created 32 mixture samples using known RNA proportions isolated from four pure cell lines (Methods). In this experiment, we employed CDSeq to estimate RNA proportions instead of cell proportions because we mixed RNA, not whole cells.

With these data, CDSeq and CIBERSORT both provided accurate estimates of sample-specific cell-type proportions. The total RMSE of CDSeq was 17% smaller than that of CIBERSORT, reflecting smaller cell-type-specific RMSEs for three out of four individual cell types (**Figure 3** and **Supplementary Figure 2**). For gene expression levels, both CDSeq and csSAM performed similarly, though CDSeq had uniformly lower RMSE across the 4 cell types (**Figure 3** and **Supplementary Figure 2**). CDSeq also outperformed csSAM by having 16% lower total RMSE for gene expression levels.

**Figure 3.**
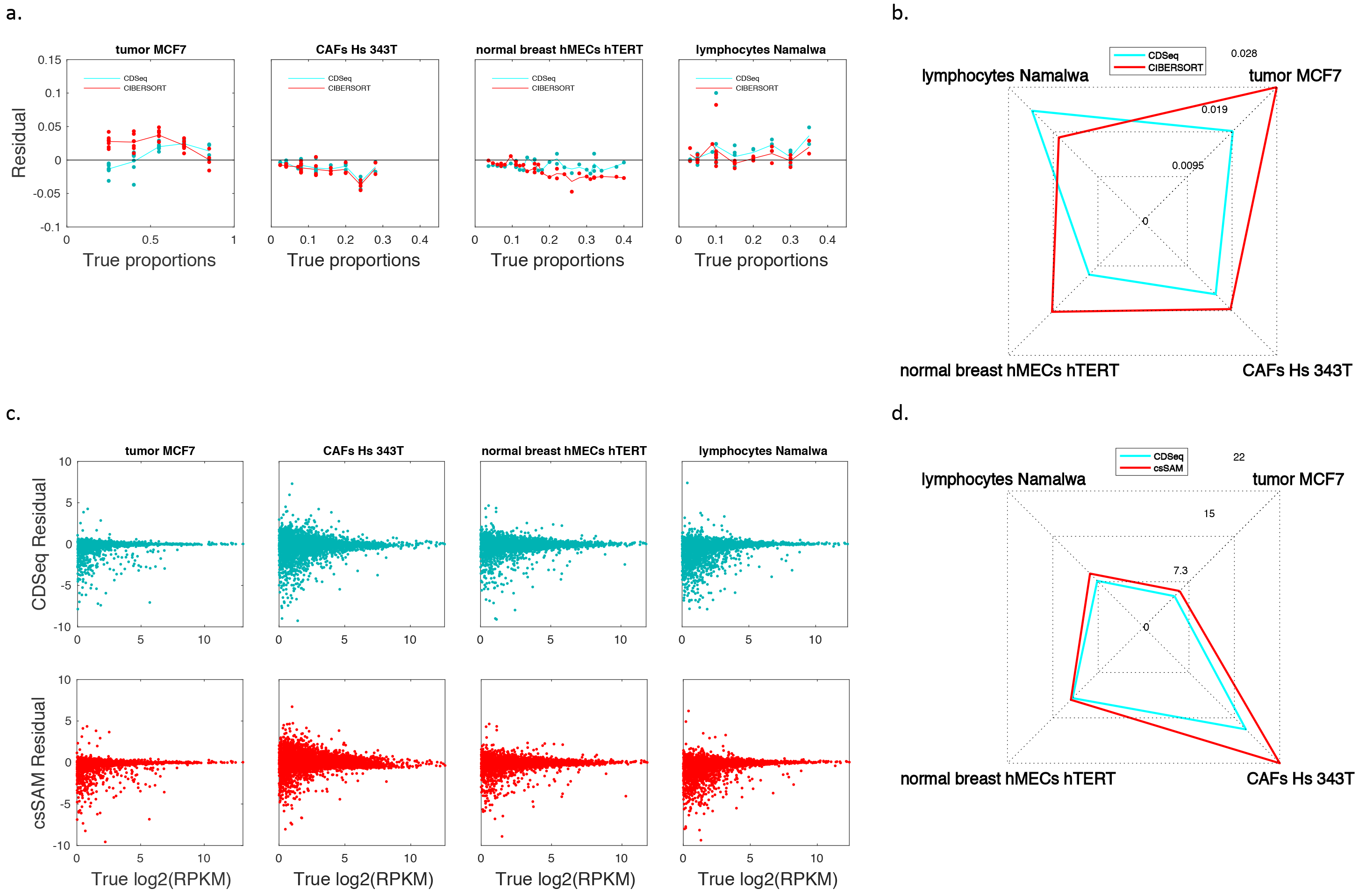
Deconvolution of mixed RNA from cultured cell lines. We cultured four pure cell lines, extracted RNA from each, and mixed the cell-type-specific RNA to create distinct 32 mixed samples with known cell-type proportions (**Supplementary Table 2**). We ran: CDseq with 4 cell types, α = 5, β = 0.5 and 700 MCMC runs; CIBERSORT with the default settings to estimate proportions while providing pure cell line expression profiles as input; and csSAM with the default settings to estimate cell-type-specific GEPs while providing true sample-specific cell-type proportions as input. (a) Difference (“residual”) between estimated and true cell-type proportion plotted against true proportion for CDSeq (green) and CIBERSORT (red). Each plotted point represents the value for a single sample. Lowess smooth was added to aid comparison. (b) Radar plot of RMSE for estimates of sample-specific cell-type proportions. CDSeq (green); CIBERSORT (red). Total RMSE summing over cell types is 17% smaller for CDseq compared to CIBERSORT. (c) Difference (“residual”) between estimated and true log_2_ gene expression level [(log_2_(RPMK)] plotted against true log_2_ gene expression level for CDseq (green) and csSAM (red). Each plotted point displays the expression value of a single gene, 19653 genes in total. (d) Radar plot of RMSE for gene expression levels. CDSeq (green); csSAM (red). Total RMSE of gene expression (summing over cell types) is 16% smaller for CDseq compared to csSAM.

### Dissecting mixtures of liver, lung, and brain cells

We next evaluated CDSeq using the experimental data set designed for csSAM^1^. The microarray data set consists of 11 mixtures (each with 3 replicates) of liver, brain and lung cells with varying known RNA proportions and included GEPs of pure liver, brain and lung. We compared the cell-type-specific GEPs estimation and the sample-specific cell-type proportion estimation of CDSeq to those of csSAM and CIBERSORT: the RMSE of CDSeq estimation is 44% lower than that of CIBERSORT; Cell-type-specific GEPs estimation provided by CDSeq and csSAM are quite close where csSAM has 1% lower RMSE than CDSeq. A detailed description of the results is given in **Figure 4** and **Supplementary Figure 3**.

**Figure 4.**
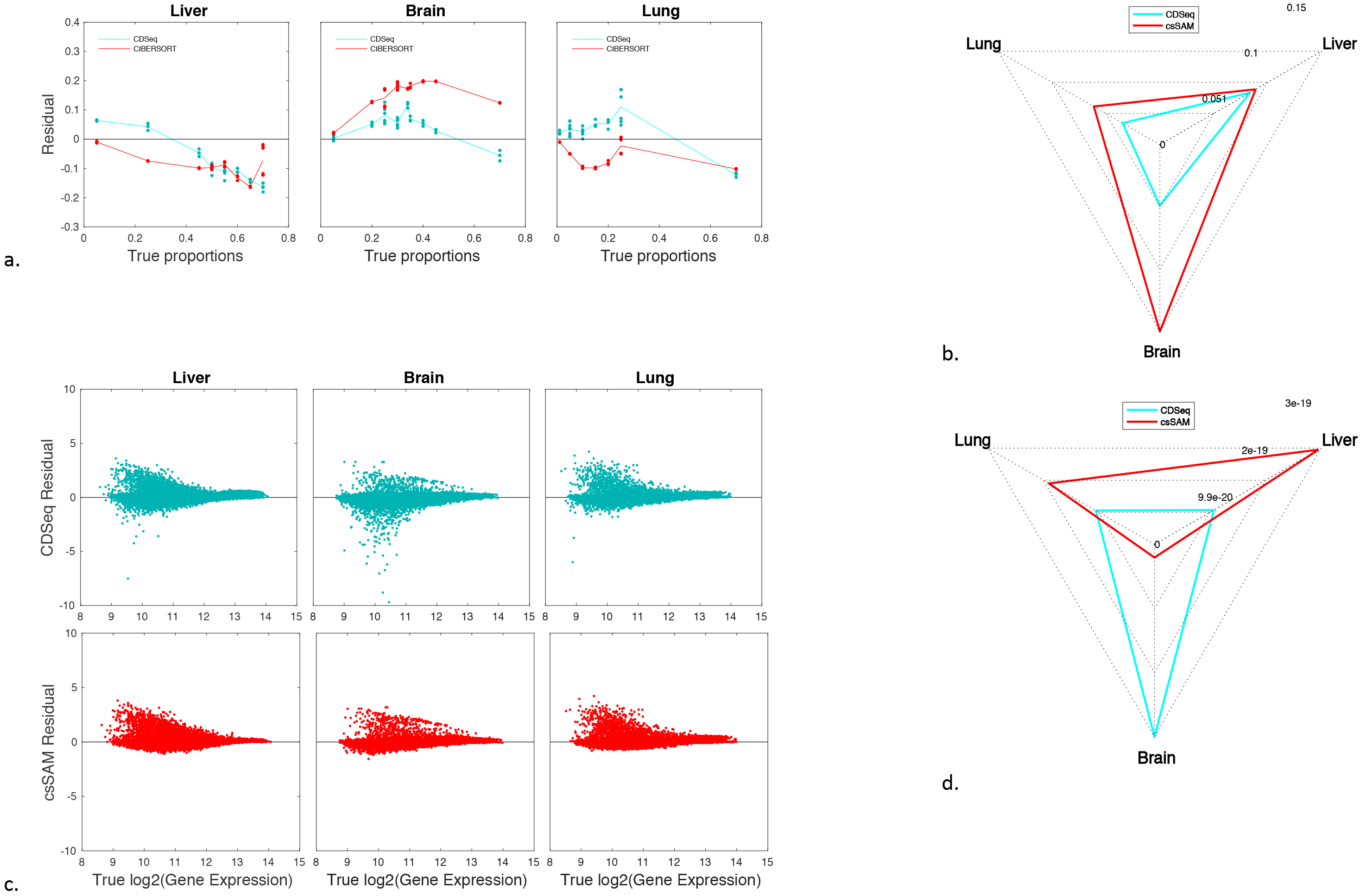
Deconvolution of mixtures of liver, brain and lung cells. The microarray data consist of 33 mixture samples with known proportions and GEPs for the three constituent pure cell lines. We ran: CDSeq by specifying three cell types, *α = 10, β = 5*, and 700 MCMC runs; CIBERSORT with default settings to estimate the proportions while providing pure cell line expression profiles as input; and csSAM with default settings to estimate cell-type-specific GEPs while providing true sample-specific cell-type proportions as input: (a) Difference (“residual”) between estimated and true cell-type proportion plotted against true proportion for CDSeq (green) and CIBERSORT (red). Each plotted point represents the value for a single sample. Lowess smooth added to aid comparison. (b) Radar plot of RMSE for estimates of sample-specific cell-type proportions. CDSeq (green); CIBERSORT (red). Total RMSE summing over cell types is 44% smaller for CDSeq compared to CIBERSORT. (c) Difference (“residual”) between estimated and true gene expression level (log_2_(gene expression) where sum of gene expression levels across all genes equals to 1 and scaled to 10^8^) plotted against true gene expression level for CDSeq (green) and csSAM (red). Each plotted point displays the expression value of a single gene, 31099 genes in total. (d) Radar plot of RMSE for gene expression levels. CDSeq (green); csSAM (red). Total RMSE of gene expression (summing over cell types) is 1% larger for CDSeq compared to csSAM.

### Evaluation using leukocyte subtypes

To test the performance of CDSeq on some extreme cases, we applied CDSeq to a set of GEPs from pure cell lines. We chose LM22^23^, designed by Newman et al.^23^, which comprises 22 human hematopoietic cell phenotypes including seven T-cell types, naïve and memory B cells, plasma cells, natural killer (NK) cells and myeloid subsets. Accordingly, we have 22 samples, each regarded as from a pure cell type; consequently, when running CDSeq, we set the number of cell types to be 22 exactly.

CDSeq identified most of the 22 samples as nearly pure examples of a single cell type, with minimal mixing except for closely related types (**Figure 5** and **Supplementary Figure 5**). When CDSeq reported nominally pure samples as mixed, the confusion was between closely related cell types; for example, naïve B cells were estimated as a mixture of naïve and memory B cells (and vice versa); and activated MAST cells were estimated as a mixture of activated and resting MAST cells (**Figure 5a**). We also ran CIBERSORT on this data set by providing LM22 itself as reference profiles (**Figure 5** and **Supplementary Figure 5**). The CIBERSORT recovered the true cell-type proportions even more accurately than did CDSeq, possibly owing to the simplicity of the problem where the inputs of mixtures and reference GEPs were identical.

**Figure 5.**
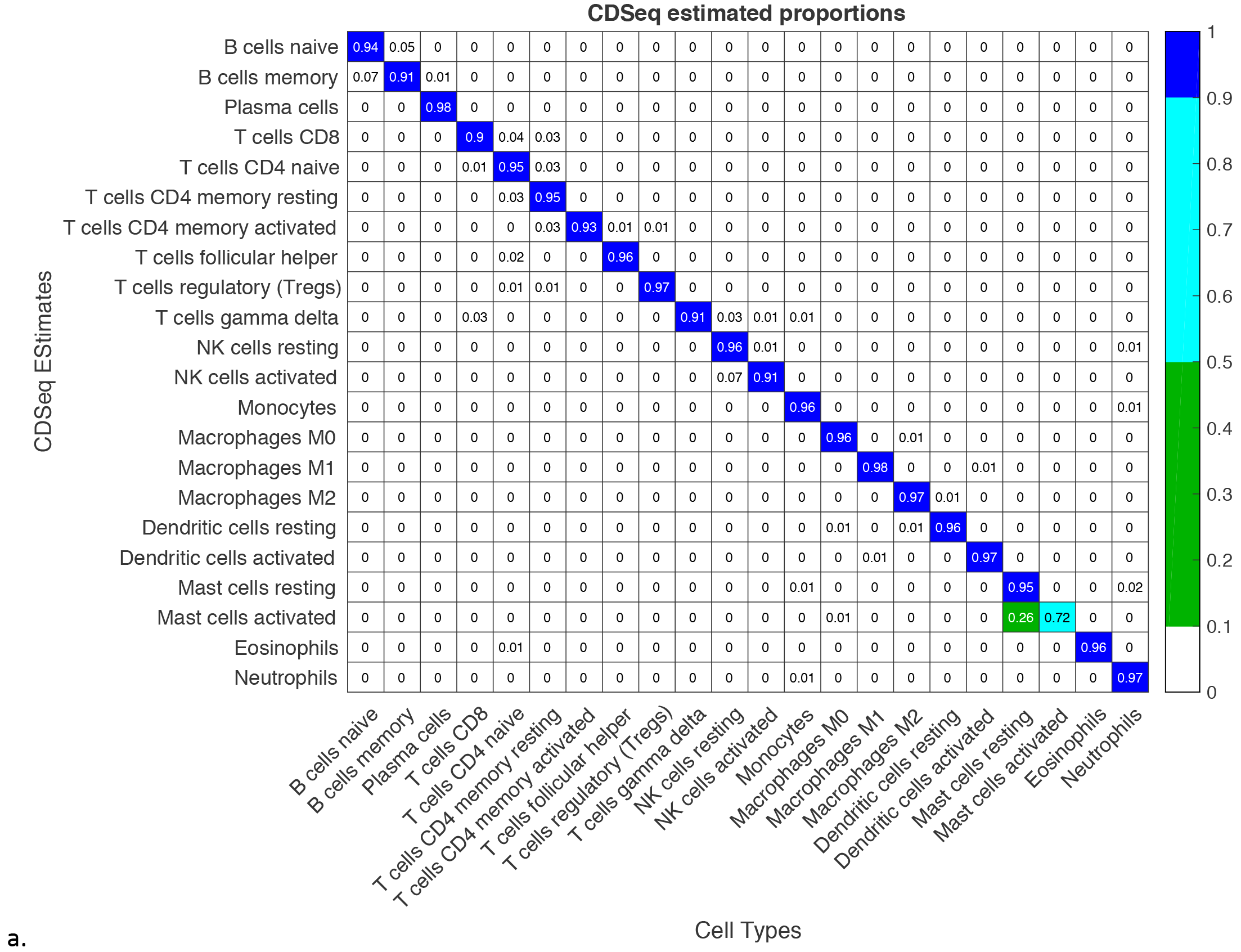
Deconvolution of LM22 data set.^23^ We ran: CDSeq with 22 cell types, α = 5, β = and 700 MCMC runs; CIBERSORT with the default settings to estimate proportions while providing pure cell line expression profiles (i.e. LM22) as input. (a) the heatmap of proportion estimation of 22 cell types by CDSeq; (b) the heatmap of proportion estimation of 22 cell types by CIBERSORT; (c) the heatmap of correlations between CDSeq-estimated GEPs and true GEPs (heatmap of correlations of true GEPs is given in **Supplementary Figure 4**); and, (d) Radar plot shows the RMSE of CDSeq (green) and CIBERSORT (red). Total RMSE is 0.33 for CDSeq and 0.2 for CIBERSORT. Notice that in figures (a) and (b) the sums of proportions along the rows are not always equal to 1 owing to the fact that we rounded off all the proportions to 2 decimal places for better visual presentation and some precisions are lost.

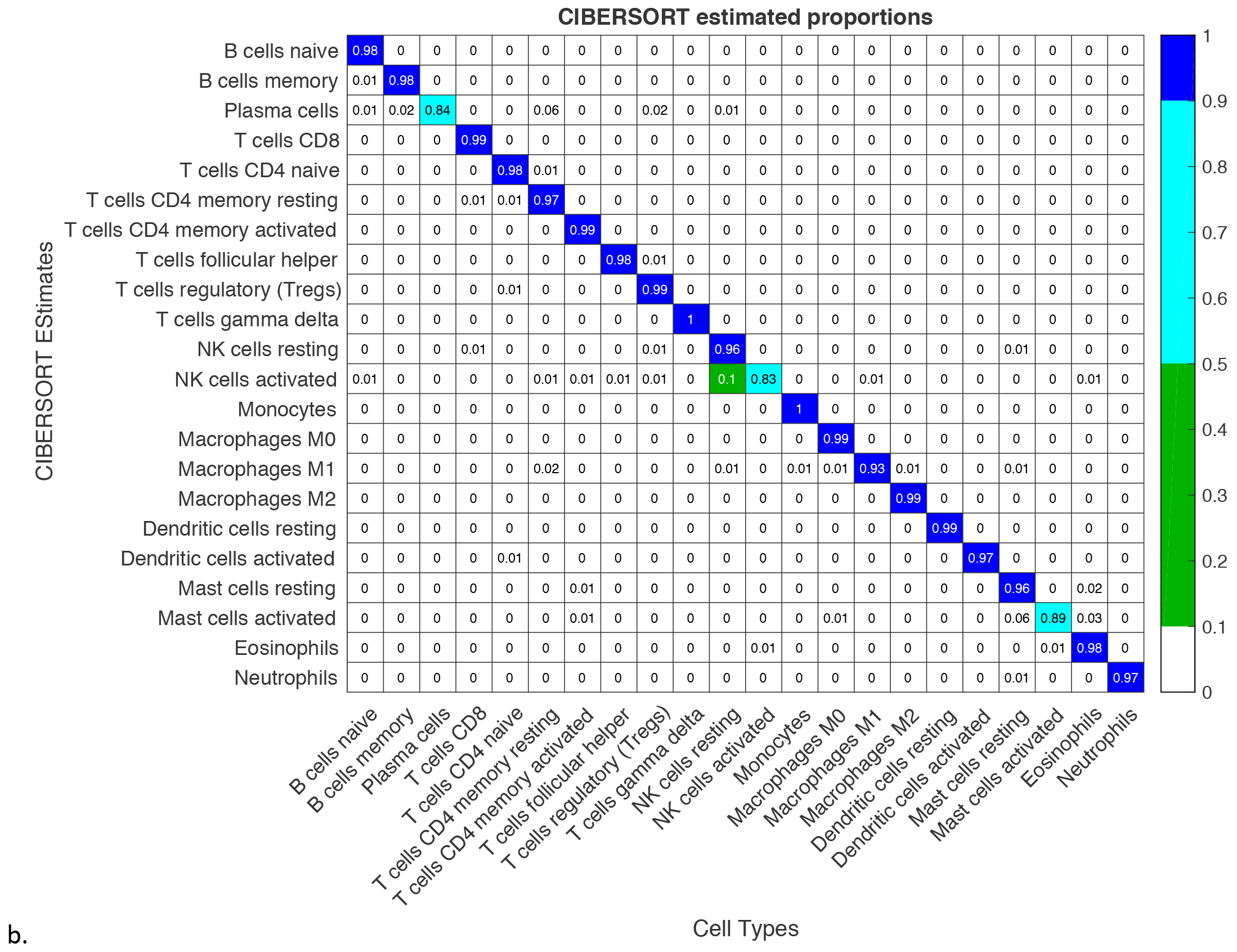

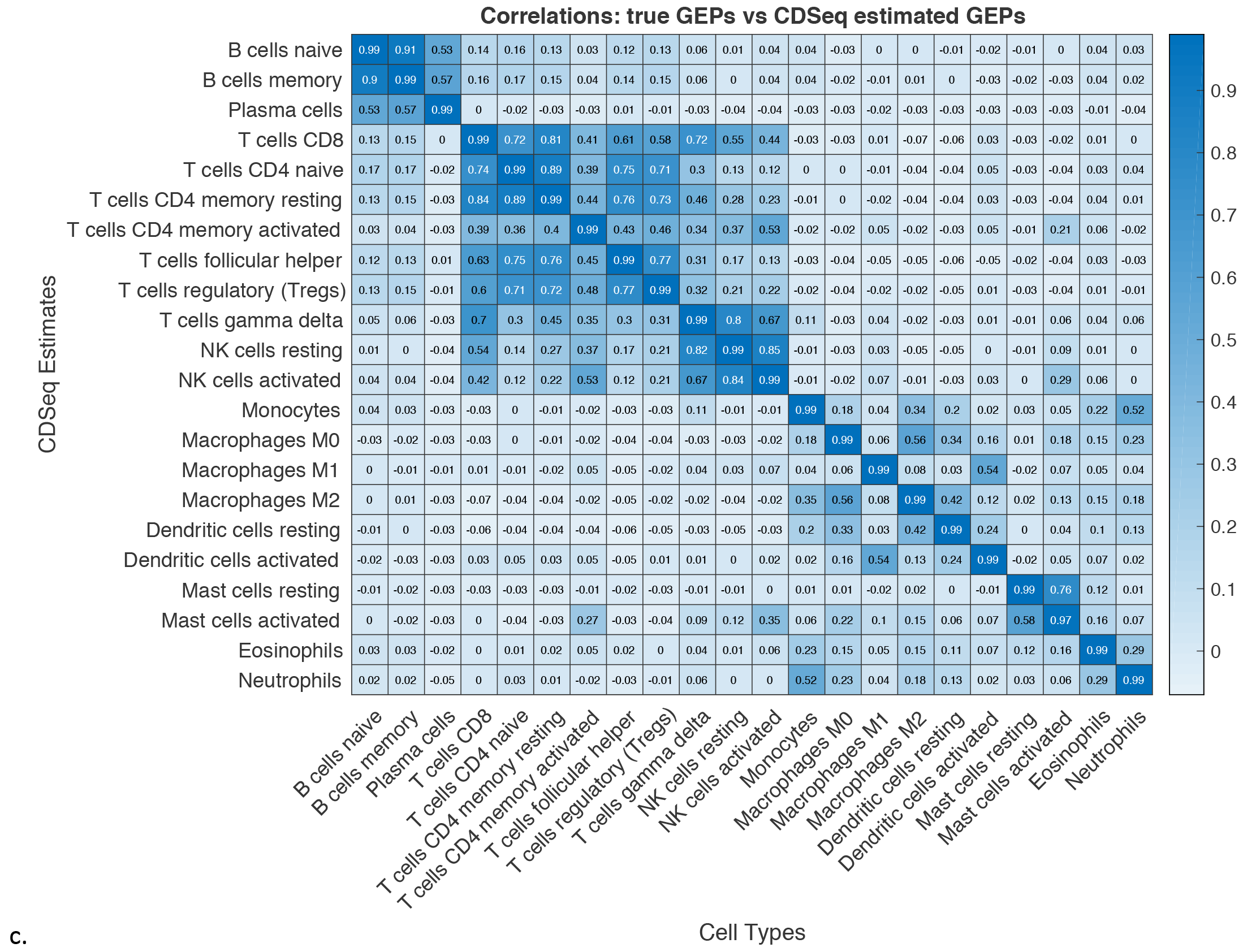

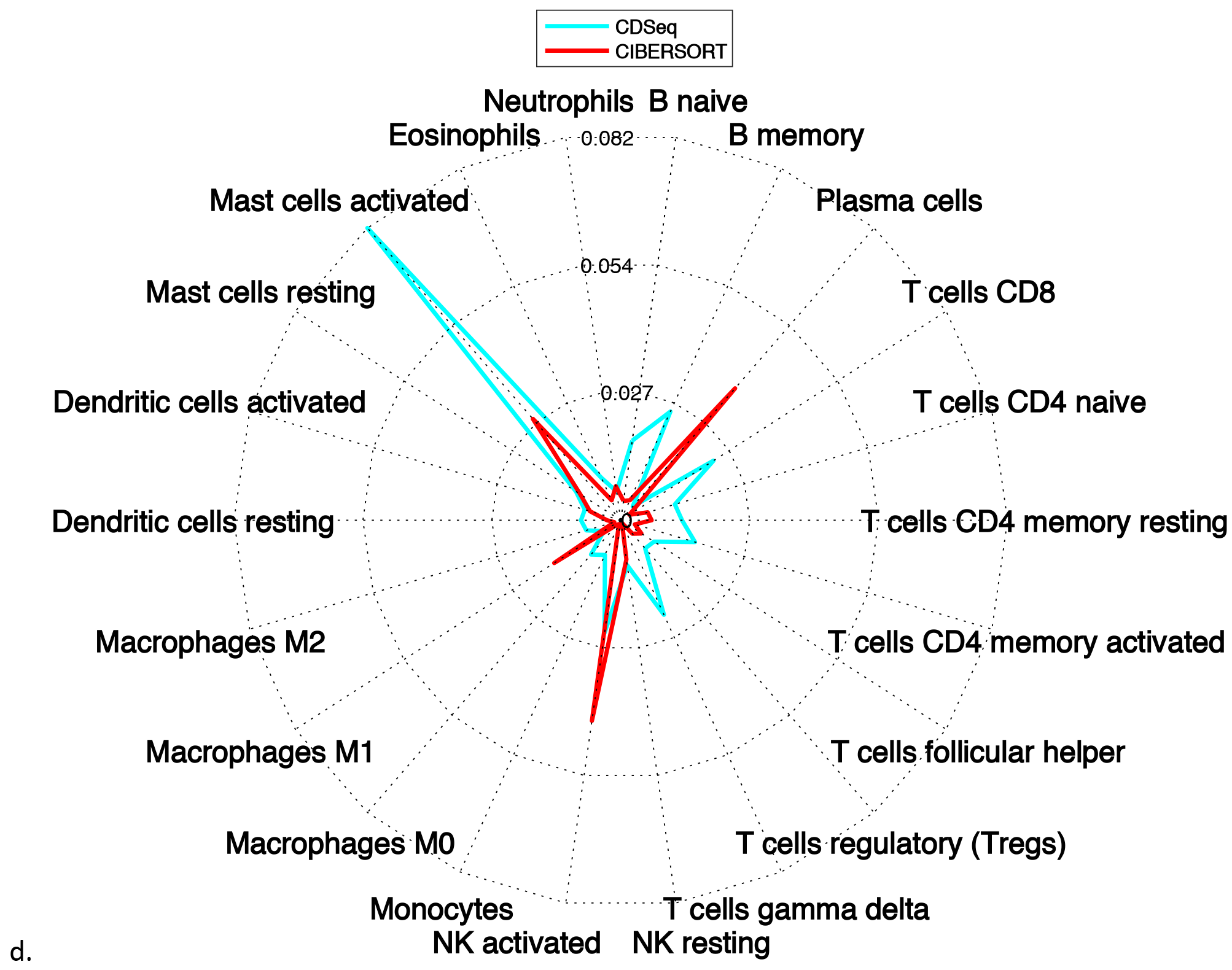

Since this data set contained only pure cell lines, we did not compare to csSAM; when the input proportions for all cell types are either one or zero as in this example, csSAM will invariably return the true GEP for each cell type.

### Immune cell analysis of lymphoma data with comparison to flow cytometry

We evaluated CDSeq against flow-cytometry measurements of leukocyte content in solid tumors. Data comprised GEPs from 14 bulk follicular lymphoma samples and corresponding flow-cytometry measurements^23^. Our goal is to estimate the proportions of B cells (naïve B cell and memory B cell) and T cells (CD8 T cell, CD4 naïve T cell, CD4 memory resting T cell, CD4 memory activated T cell, follicular helper T cell, regulatory T cell) in those 14 samples using CDSeq. We set the number of cell types to be eight (the number of all B cell and T cell subtypes in our reference file).

To match the anonymous cell types identified by CDSeq to actual B cell or T cell subtypes, we compared the GEPs estimated by CDSeq with the GEPs of the eight relevant B and T cell subtypes from LM22 using correlation. We considered an anonymous CDSeq-identified cell type to match one of the B cell or T cell subtypes if the Pearson correlation of their GEPs exceeded 0.6. (Thus, a CDSeq-identified cell type could match none or multiple leukocyte subtypes and vice versa.) We found that three CDSeq-identified cell types (cell type 1, 3, 7 in **Supplementary Figure 6**) matched no leukocyte subtypes, one CDSeq-identified cell type (cell type 5 in **Supplementary Figure 6**) matched only memory B cells, and four CDSeq-identified subtypes (cell type 2, 4, 6, 8 in **Supplementary Figure 6**) matched both naïve and memory B cells. No CDSeq-identified cell types matched any T cell subtypes (**Supplementary Figure 6**). Thus, CDSeq was not fully successful at resolving subtypes of either B cells or T cells: CDSeq-identified cell types could be matched to B cells, the majority cell type in the lymphoma samples determined by flow cytometry, but could not be matched to the lower abundance T cells.

This lack of success with low abundance cell types could have several sources. First, because of differences in tumor microenvironment across samples, a cell type embedded in a solid tumor may exhibit quite a different GEP from sample to sample, and a different GEP than it might exhibit in pure culture. Second, CDSeq estimates may represent some average effects of the underlying signals, mathematically, the output may correspond to local optima owing to a complex posterior distribution of the parameters. To overcome this difficulty with low abundance cell types, we propose augmenting the input data from mixed samples with GEPs from pure lines of the cell types likely to be in the mixed samples. We call this strategy a quasi-unsupervised learning strategy (Methods). The idea is, without introducing labels, to provide the algorithm some guidance in searching for cell-type-specific signals in solid tumors. In this example, we appended 8 B cells and T cells lymphocyte GEPs from LM22 to the bulk measurements as input to CDSeq.

To see whether the quasi-unsupervised strategy could improve detection of T cell subtypes, we compared the estimated GEPs from quasi-unsupervised CDSeq to the reference GEPs of eight B cell and T cell subtypes via correlations as before. With the quasi-unsupervised strategy and the 0.6 correlation threshold, performance of CDSeq improved for T cell subtypes (**Figure 6**). Two CDSeq-identified cell types (cell type 6 and 8 in **Figure 6**) matched either all six or five of six T cell subtypes but no B cell subtypes; five CDSeq-identified cell types (cell type 1, 3, 4, 5 and 7 in **Figure 6**) matched one or both B cell subtypes but no T cell subtypes; and one CDSeq-identified cell type (cell type 2) matched none of the eight lymphocyte subtypes. Nevertheless, CDSeq still could not fully resolve subtypes among B cells and T cells. Consequently, we decided to estimate relative proportions of B cells and T cells in each sample regardless of subtype by summing the relevant subtype proportions while ignoring the estimated proportions of unmatched estimated cell type 2, then renormalizing the relative proportions to sum to one. The resulting estimated relative proportions were statistically significantly correlated with corresponding flow cytometry measurements (p < 0.05) (**Figure 6**).

**Figure 6.**
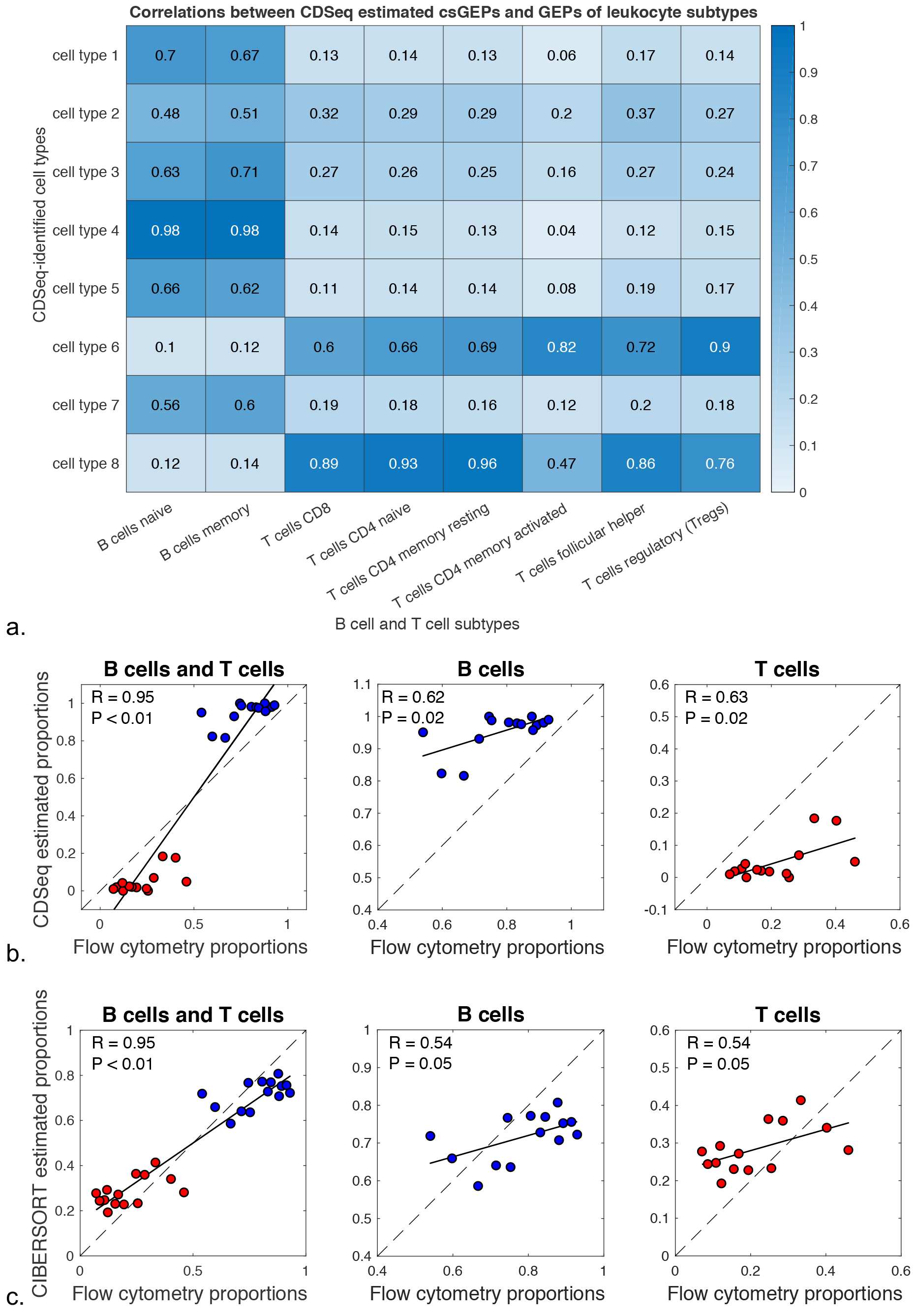
Deconvolution of B cells and T cells in lymphoma. We applied CDSeq on lymphoma data set^23^ employing the quasi-unsupervised strategy (Methods) by appending the B cells and T cells GEPs to the GEPs of 14 solid tumor samples. We ran CDSeq with the number of cell types set at 8, *α* = 0.5, *β* = 0.5. In (b) and (c), the black solid line is the linear regression line; the black dashed line is the x=y line, R is the correlation coefficient, and P is the p-value for testing the null hypothesis of no correlation. (a) Correlations between CDSeq-estimated cell-type-specific GEPs and GEPs for B cell and T cell subtypes provided by LM22; and (b) CDSeq-estimated proportions of B cells (blue dots) and T cells (red dots): the first panel displays the estimated relative proportions for both B cells and T cells, the second and third panels present B cells and T cells separately; and (c) CIBERSORT-estimated relative proportions of B cells (blue dots) and T cells (red dots) with the three panels providing analogous information to those in Figure 6b.

We did not apply csSAM since we had no ground truth for cell-type-specific GEPs available for accessing the accuracy and comparison between csSAM and CDSeq.

### CDSeq on deep deconvolution

Deep deconvolution refers to the problem of using a whole blood or PBMC sample to estimate the proportions and expression profiles of a greater number of cell subtypes, going further down into the hematopoietic tree^10^. To assess CDSeq’s performance on deep deconvolution, we used a set of 20 PBMC samples^23^. To evaluate performance, we also used information provided in the LM22 dataset^23^: namely, flow-cytometry measurements for nine of the 22 leukocyte sub-types (the only sub-types with flow cytometry available). That LM22-provided GEPs of about half of these nine sub-types were highly collinear (**Supplementary Figure 4**) should challenge CDSeq’s ability both to find the corresponding GEPs of those nine sub-types in the 20 PMBC samples and to accurately estimate their proportions. We first ran CDSeq in fully unsupervised mode and set the number of cell types to be 22. In comparing the estimated GEPs to the GEPs of LM22, we found that CDSeq could not uncover the nine sub-types (**Supplementary Figure 7**), possibly because collinearity of GEPs among sub-types.

To improve estimation by reducing complexity, we turned to the quasi-unsupervised strategy when running CDSeq by appending the 22 GEPs of LM22 to the 20 samples, 42 samples in total. Using the 0.6 correlation threshold to match CDSeq-identified cell types to the corresponding 22 leukocyte sub-types, we found that the quasi-unsupervised strategy improved CDSeq’s performance (**Figure 7**): one CDSeq-identified cell type matched both naïve and activated B cells; another matched both resting and activated mast cells; two CDSeq-identified cell types did not match any of the 22 LM22 known sub-types; the remainder matched only one LM22 sub-type each. When comparing CDSeq-estimated cell-type relative proportions to flow-cytometry measurements for the nine sub-types, we combined the proportions of naïve B cells and memory B cells as one overall B cell sub-type because the estimated GEPs of these two were highly correlated (**Figure 7** and **Supplementary Figure 7**). In restricting attention to the resulting eight sub-types, we renormalized their proportions to sum to one for comparison with corresponding flow cytometry measured proportions. For six of the eight sub-types, the CDSeq-estimated relative proportions were significantly correlated (p<0.05) with the flow-cytometry-based relative proportions. The correlations with activated memory CD4 T cells and *γδ* T cells were not significant (p=0.31 and 0.07, respectively). The CIBERSORT estimated relative proportions were significant correlated (p<0.05) with the corresponding flow-cytometry-based relative proportions for all sub-types except *γδ* T cells (p=0.19). In an overall comparison of CDSeq and CIBERSORT estimates, however, the total RMSE of CDSeq was about 6% lower than that of CIBERSORT. On the other hand, the estimated relative proportions by both CDSeq and CIBERSORT showed systematic bias in departing from equality with the flow-cytometry-based proportions. Besides the possible technical issues of flow cytometry and the fidelity of the LM22 reference profiles, another possible reason for this systematic bias with this microarray data is that flow cytometry reports relative cell proportions whereas CDSeq and CIBERSORT report relative RNA proportions. Though CDSeq is capable of reporting either RNA proportions or cell proportions from RNA-seq raw counts, it can report only RNA proportions with microarray data.

**Figure 7.**
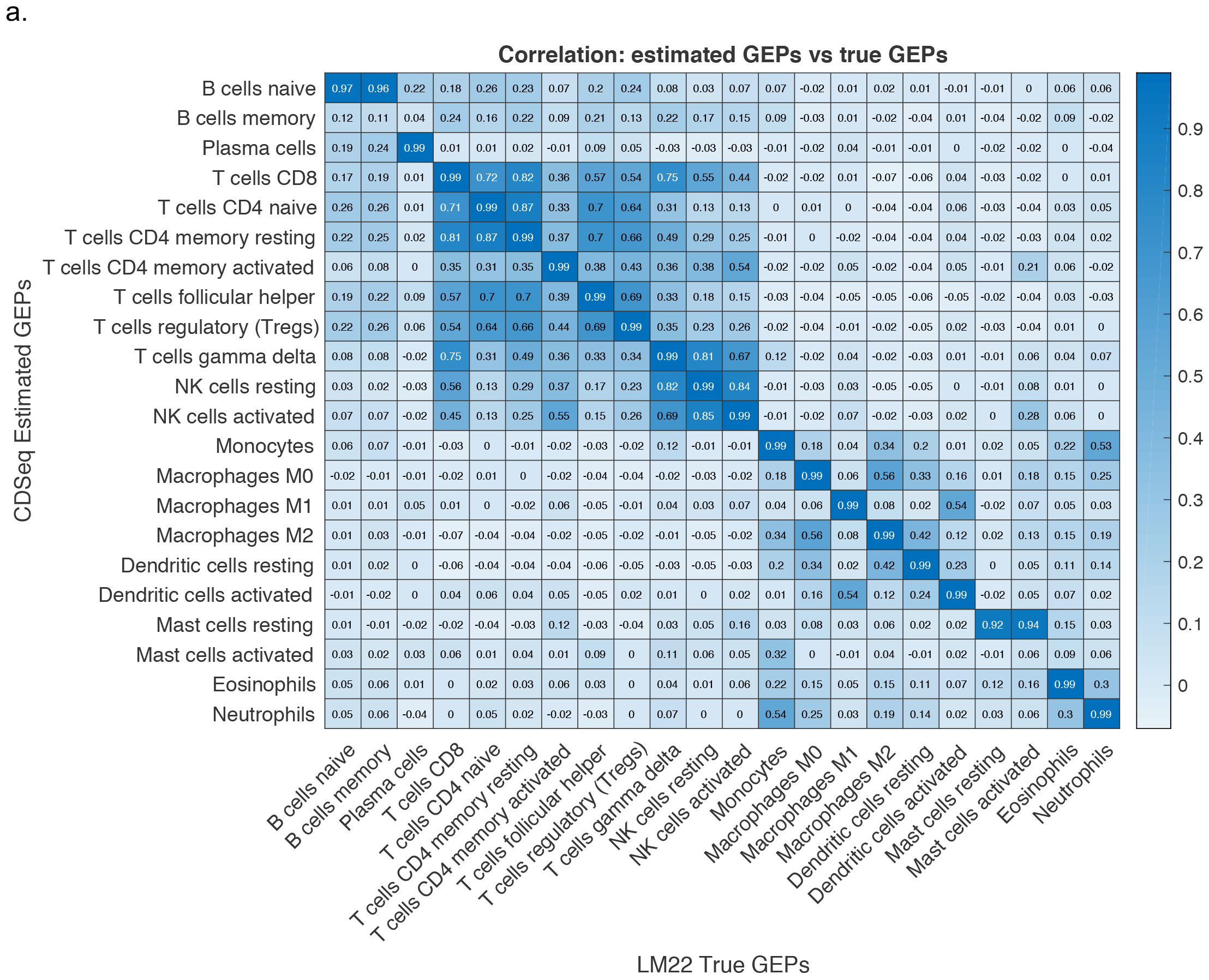
Deep deconvolution of PBMC data. We applied CDSeq using quasi-supervised learning strategy to the 20-sample PBMC data set after appending 22 LM22 GEPs, 42 samples in all. We ran CDSeq with 22 cell types, *α* = 50, *β* = 20. In (b), the black line is the linear regression line; the dashed line is the x=y line; R is the correlation coefficient; and P is the p-values for testing the null hypothesis of no correlation: (a) Correlations between CDSeq estimated cell-type-specific GEPs and LM22 GEPs (heatmap of correlations of true GEPs is given in **Supplementary Figure 4**); (b) Upper panel CDSeq-estimated proportions for eight cell types (green dots); lower panel is CIBERSORT-estimated proportions for eight cell types (red dots); and, (c) Radar plot for RMSE of CDSeq (green) and CIBERSORT (red).

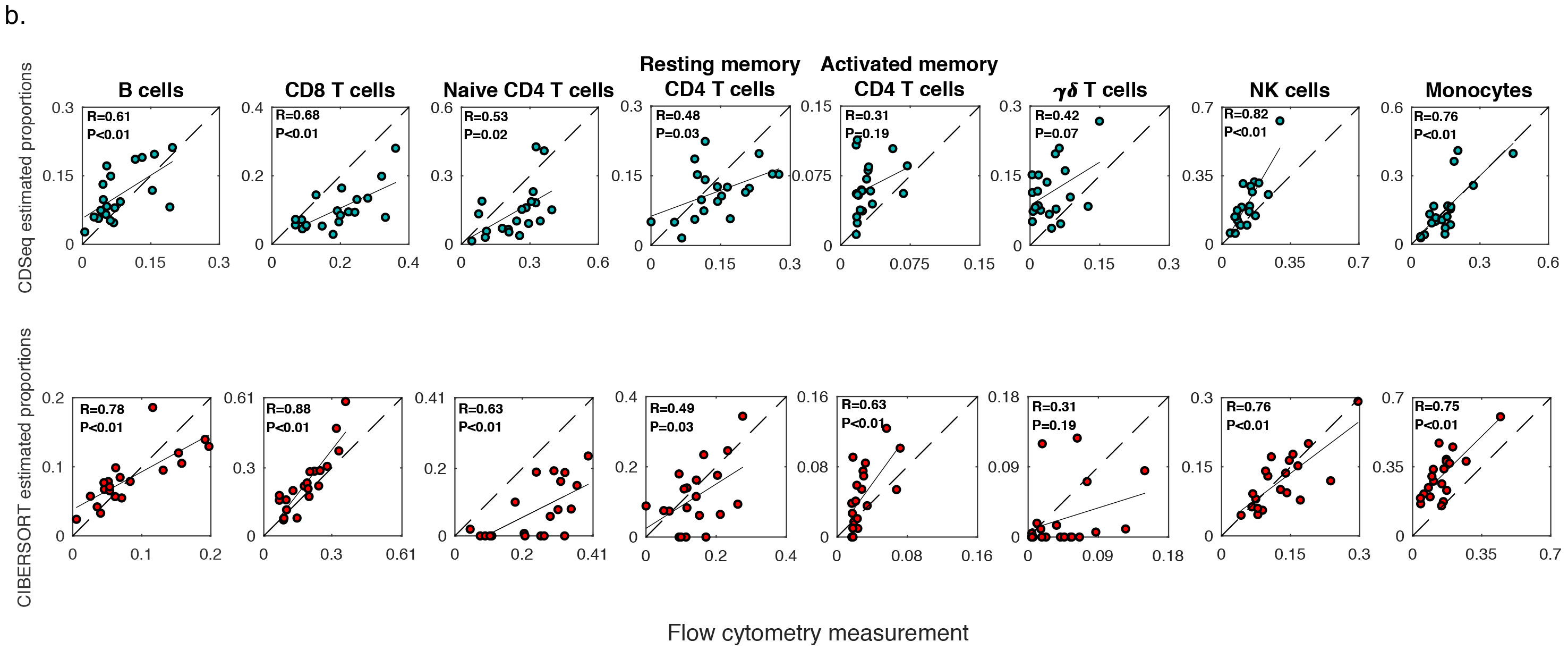

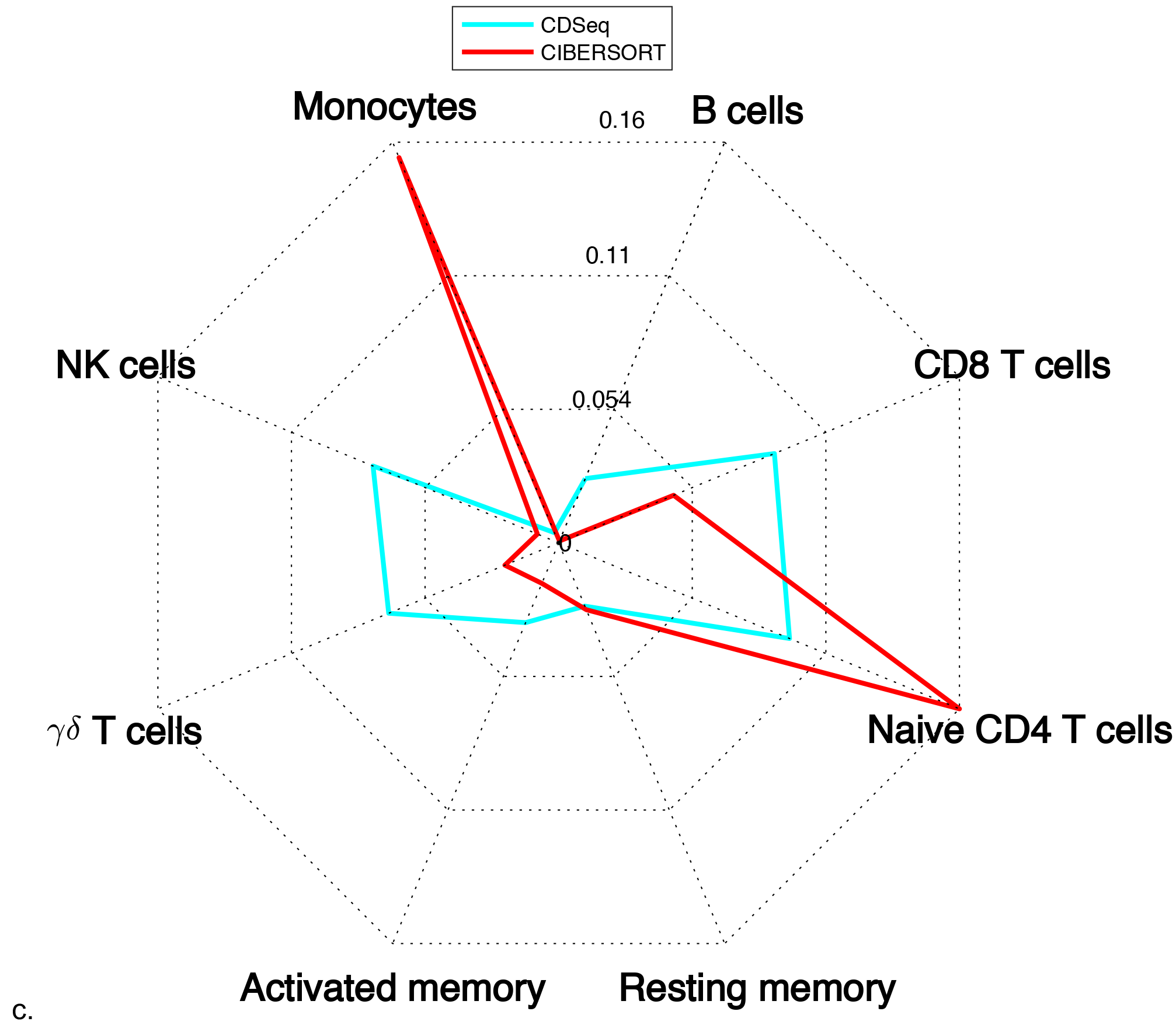

### Estimating the number of cell types present from the data

We have been applying CDSeq by fixing the number of cell types at the correct number, since we know it in advance. CDSeq can, however, estimate the number of constituent cell types in a collection of samples, if necessary, by maximizing the posterior distribution (Methods). The framework of CDSeq is built for RNA-seq raw count data, therefore, raw count data is required for estimating the number of cell types. Consequently, we did not apply this feature for microarray data.

Applying this method to the synthetic data and to the data on mixed RNA described above correctly estimated number of cell types in each case (**Figure 8**).

**Figure 8.**
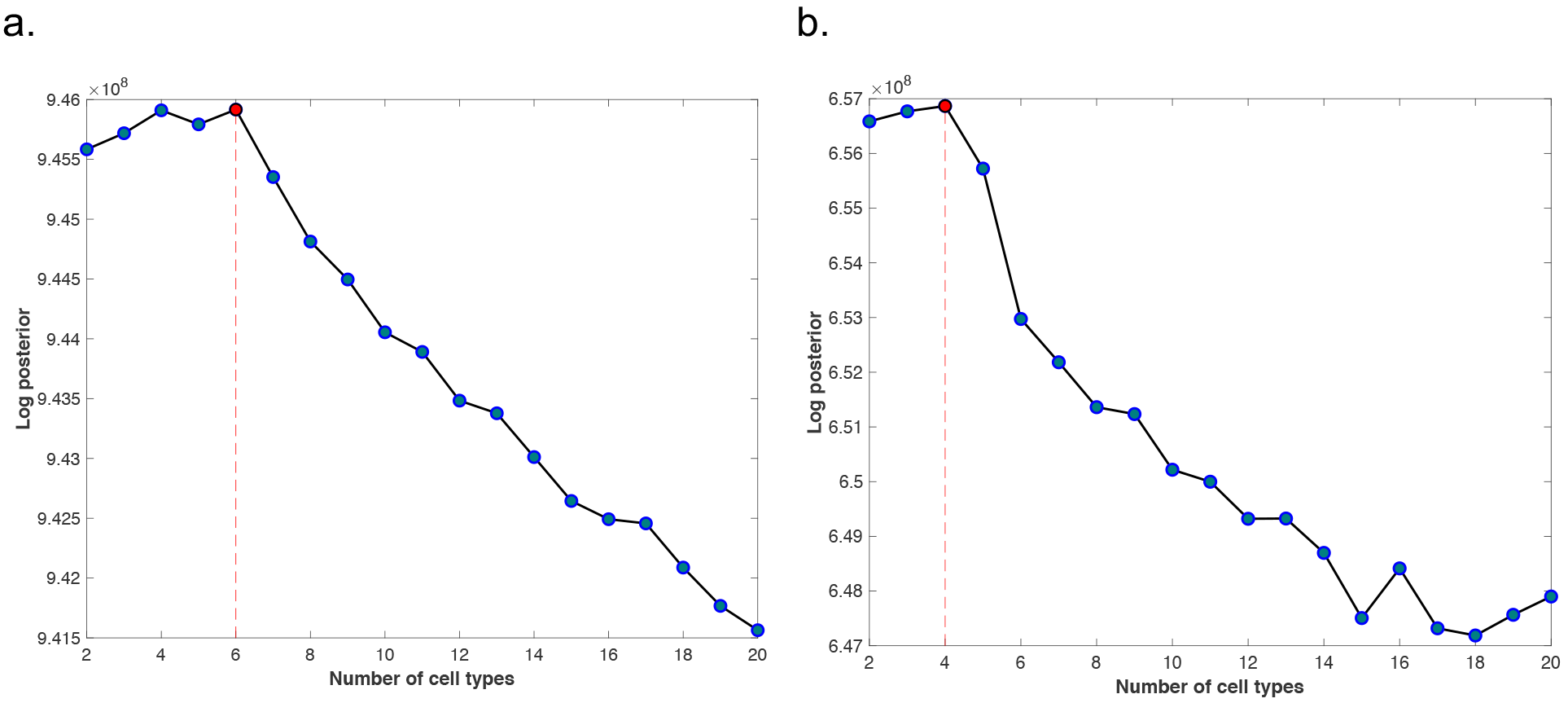
Estimating the number of cell types. The maximum of the log posterior provides an estimate of the number of cell types. (a) synthetic data; (b) mixed RNA data. In each data set, the method estimated the true number of cell types correctly.

## Discussion

We present CDSeq for deciphering heterogeneity in RNA-seq data measured on biological samples. CDSeq, as a complete deconvolution method, exhibits substantial advantages over existing deconvolution methods, such as csSAM^1^ and CIBERSORT^23^, as it only requires expression data from mixtures and outputs estimates of both cell-type-specific GEPs and sample-specific cell-type proportions. In addition, we argue that our probabilistic model is of conceptual advance over methods using matrix decomposition^15,18^ or regression techniques.^1,23^ First, our generative model explicitly considers how reads are generated and is able to estimate cell proportions instead of RNA proportions whereas matrix decomposition or regression based methods are not. Second, our model employs multinomial random variable to capture the stochastic nature of reads and therefore inherently imposes the constraint that proportions are nonnegative and sum to one on the parameters of interest; whereas matrix decomposition or regression-based methods need to impose additional constraints on the parameter space that can impose additional technical challenges for numerical procedures.

We assessed CDSeq using both synthetic and real experimental data and saw generally good estimation performance. Often complete deconvolution by CDSeq was as or more accurate than partial deconvolution by CIBERSORT to estimate cell-type proportions or by csSAM to estimate GEPs.

CDSeq, an unsupervised data mining tool, is fully data-driven and allows simultaneous estimation of both cell-type-specific GEPs and sample-specific cell mixing proportions. In some real data analyses when constituent cell types had highly correlated GEPs, the cell types found by CDSeq lacked a one-to-one correspondence with the known component cell lines. To ameliorate this problem, we proposed a quasi-unsupervised approach. It involves augmenting the available GEPs from heterogeneous samples with GEPs from pure cultures of the cell types anticipated to be constituents. We showed that this quasi-unsupervised approach can improve CDSeq’s performance in lymphoma and deep deconvolution examples.

A limitation of current CDSeq model is the impossibility of fine tuning the hyperparameters to obtain optimal results without ground truth. In practice, we suggest setting *α* = 5, *β* = 0.5. When heterogenous samples are likely dominated by one or two cell types, setting *α* < 1 may help; when cell-type-specific GEPs are likely to have relatively high correlation, setting *β* > 1 may help – though we cannot specify a definitive threshold for high correlation. From a practical point of view, the higher the correlations are, the fuzzier discovered signal would be. Another potentially helpful technique is the quasi-unsupervised strategy. Efforts at enabling CDSeq to self-adjust hyperparameter based on given data are underway. Another possible extension for current model is that the fundamental multinomial model used for gene expression imposes a certain negative correlation between expression counts at different loci. However, it is conceivable that, because genetic pathways can be regulated as units, the counts could be positively correlated among certain subsets of genes. The current CDSeq model cannot handle that kind of correlation structure. In addition, we suggest excluding unnecessary genes (depending on the application) to reduce the running time of CDSeq by consuming less memory.

We expect that CDSeq will prove valuable for analysis of cellular heterogeneity on bulk RNA-seq data. It provides a practical and promising alternative to methods that require expensive laboratory apparatus and extensive labor yet entail possible loss of a system perspective by isolating individual cells from heterogeneous samples. Application of CDSeq will aid in deciphering complex genomic data from heterogenous tissues.

## Methods

### Synthetic mixtures

We generated synthetic gene expression profiles for 40 synthetic mixture samples using gene expression profiles for six pure cell lines. We downloaded expression profiles from the CSHL website for: normal fetal lung fibroblast, normal B-lymphocyte (blood), normal mammary epithelial cells (mammary gland), normal umbilical vein endothelial cells (blood vessel), breast epithelial carcinoma, and normal CD14-positive cells from human leukapheresis production. To artificially amplify the confounding factor of differences in cell-type-specific RNA quantity, we multiplied the cell line reads count by a predefined vector to rescale the RNA amounts. Specifically, we randomly chose to double RNA-seq reads count of the normal B lymphocyte and normal mammary epithelial breast cell lines. We then randomly generated mixing proportions (**Supplementary Table 1**) that specified the proportion of cells of each type in each synthetic sample using a Dirichlet distribution with a parameter vector having all six entries equal to 5.

### Experimental mixtures and gene expression profiling

MCF7 cells were obtained from Duke University Cell culture facility and cultured in DMEM medium supplemented with 10% fetal bovine serum (FBS). Namalwa cells were a gift from Dr. Sandeep Dave, Duke University, and were cultured in RPMI medium supplemented with 10% FBS. Hs343T and hTERT-HME1 (ATCC) were cultured in HuMEC Ready medium (Thermo Fisher Scientific).

In brief, total mRNA was prepared from Namalwa (Burkitt’s lymphoma), Hs343T (fibroblast line derived from a mammary gland adenocarcinoma), hTERT-HME1 (normal mammary epithelial cells immortalized with hTERT), and MCF7 (estrogen receptor positive breast cancer cell line). mRNA samples were diluted to 100 ng/*μ*l and mixed in different proportions (**Supplementary Table 2**). Global mRNA abundance of the four pure cell lines and of the mixed RNA samples was profiled by RNA-sequencing.

Sequencing libraries were prepared using TruSeq RNA sample preparation kit v2 (Illumina). 75-bp single end sequencing was performed on the NextSeq sequencer (Illumina). After obtaining the fastq data, we first ran cutadapt (version 1.12) for trimming adapter sequences. Secondly, we mapped reads to the genome using STAR (version 020201). Lastly, we used featureCounts (version 1.5.1) to generate raw read counts data as the input for our algorithm. The code for processing the fastq data using cutadapt, STAR and featureCounts are available at https://github.com/kkang7/CDSeq_011.

Based on RNA-sequencing, we found contamination in the pure cell lines by examining the gene expression of ***KRT5***. ***KRT5***, a marker specific to HME-hTERT, should be completely absent from cancer-associated fibroblasts (CAFs); however, it was present in CAF samples at about 20% of the levels found in hTERT-HME1. This contamination is possibly attributable to CAFs being derived from tumors so that they were probably contaminated with a small portion of tumor tissue. *In vitro*, this small portion of tumor cells had huge growth advantage and, thus, became significant. In summary, CAF samples were not pure but contained about 20% RNA from HME-like cells. We are not certain if the contamination comes from hTERT-HME1 cells or other cancer cells which express endogenous TERT. To alleviate this problem, we considered the proportions of CAF (given in **Supplementary Table 2**) to be 80% of CAF and 20% of HME and adjusted the proportions accordingly in the comparisons (**Figure 3**).

### Overview of CDSeq

We developed a novel statistical model which aims at extracting cell-type-specific information using bulk RNA-seq data from heterogeneous samples. The proposed method performs complete deconvolution. Using only bulk RNA-seq expression data for multiple samples as input, it provides estimates of both cell-type-specific GEPs and sample-specific cell-type proportions simultaneously, in an unsupervised fashion. Our model was inspired by latent Dirichlet allocation (LDA)^26^, a probabilistic model designed for natural language processing. LDA was designed to use text corpora as input and extract essential structure, namely, the topics that constitute the content of documents in the corpus. The problem of deriving abstract yet meaningful topics from a corpus of documents shares a fundamental similarity with the problem of extracting cell-type-specific information from bulk RNA-seq data. The original LDA model cannot, however, fully capture the complexity of bulk RNA-seq data. For example, different cell types may produce different amounts of RNA, a circumstance that may bias estimates of cell-type proportions. Our model extends LDA model in the following ways: first, the random variable that models cell-type-specific GEPs depends on gene length; second, the probability of having a read from a cell type depends on both the proportion of that cell type present in a sample and the typical amount of RNA produced by cells of that type. Some existing methods are based on the LDA model^19,20,27^; however, those methods were designed for partial deconvolution and require cell-type-specific GEPs as input.

To describe our model and the statistical inference scheme, we first introduce the notation. Let M denote the number of samples and T denote the number of cell types comprising each heterogeneous sample. We model the vector containing the cell-type-specific proportions for sample *i*, denoted *θ*_*i*_ = (*θ*_*i*,1_, *θ*_*i*,2_,…, *θ*_*i,T*_) ∈ *S*^*T*^, where *S*^*T*^ denotes a (T-1)-simplex, as a Dirichlet random variable with hyperparameter 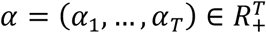. Next, let G denote the number of genes in the reference genome to which reads are mapped. We denote the GEP of pure cell type *t*, a vector of gene expression values for the entire genome normalized to sum to 1, as *ϕ*_*t*_ = (*ϕ*_*t*,1_, *ϕ*_*t*,2_,…, *ϕ*_*t,G*_) ∈ *S*^*G*^, where *S*^*G*^ denotes a (G-1)-simplex and model it as a Dirichlet random variable with hyperparameter 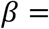 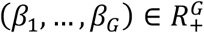. With T cell types in all M samples, the matrices *θ* = [*θ*_1_,… *θ*_*M*_] and *ϕ* = [*ϕ*_1_,…,*ϕ*_*T*_] encapsulate all the features that we seek to estimate from the data based on our model.

We denote the true GEP of heterogeneous sample *i* by ϕ_*i*_ = (*ϕ*_*i*,1_, *ϕ*_*i*,2_,…, *ϕ*_*i,G*_) ∈ *S*^*G*^. Φ_*i*_ is a weighted average of the pure cell-type GEPs with weights given by the sample-specific cell-type proportions, namely, 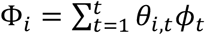. This random variable controls the rate of generating RNA copies from genes.

We do not observe the true *ϕ*_*i*_ directly but instead observe reads from each sample and the read assignments to genes. Assume that the length of every sequenced read, denoted *m*, is the same. Let categorical random variable *r*_*i,j*_ ∈ {1,…, 4^*m*^} denote read *j* from sample *i*, and let categorical random variable *g*_*i,j*_ ∈ {1,…, *G*} denote the gene or transcript assignment of read *r*_*i,j*_. Both 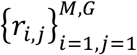 and 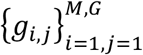 are observed for every heterogenous sample. In transcript *k*, the number of positions in which a read can start is 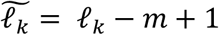 where *ℓ*_*k*_ is the length of transcript *k*. The adjusted length 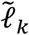 is called the effective length of transcript *k*. If the reads are mapped to genes instead of transcripts isoforms, then we need to consider the effective length of gene, denoted by 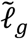, which is total length of all the transcripts comprising the gene after projection into genomic coordinates. Different cell types may generate different amounts of RNA owing to their varying sizes, therefore we employ a Poisson random variable with parameter *η*_*t*_ to model the number of reads generated from cell type t. Let *η* = (*η*_1_,…, *η*_*T*_). Parameter *η* can be estimated from RNA-seq read counts from pure cell types using the unweighted sample mean, a maximum likelihood unbiased estimator. If such information is not available, CDSeq will treat *η* as a unit vector indicating no differences in cell sizes.

Finally, to complete specification of our model, we need to be able to assign reads in the heterogenous sample to individual cell types; thus, we introduce a latent categorical random variable *c*_*i*,*j*_ ∈ {1,…, *T*} that is the cell type indicator of read *r*_*i,j*_.

Our model specifies that RNA-seq reads from bulk tissues are generated as follows:

1. Generate gene expression profiles for different cell types, i.e., *ϕ*_*t*_ ~ *Dir*(*β*) for cell type *t*,*t* = 1,…,*T*.
2. Choose *θ*_*t*_ ~ *Dir*(*α*) which denotes the mixture proportion of different cell types in the sample *i*,*i* = 1,…,*M*.
3. For each of the *N*_*i*_ RNA-seq reads in sample *i*, where *N*_*i*_ denotes the total reads of sample *i*

a. Choose a cell type *c*_*n*_ ~ *multinomial*((*θ*, *η*), where *n* = 1,…, *N*_*i*_ and the notation (*θ*, *η*) means the multinomial distribution depends on both parameters. (See Supplementary Materials on statistical inference for details.)
b. Choose a gene *c*_*n*_ ~ *p*(·|*c*_*n*_, *β*, *ϕ*) = *multinomial*(*ϕ*_*c*_*n*__)
c. Generate a read sequence 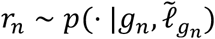 where 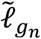 is the effective length of transcript *g*_*n*_.

**Figure 9.**
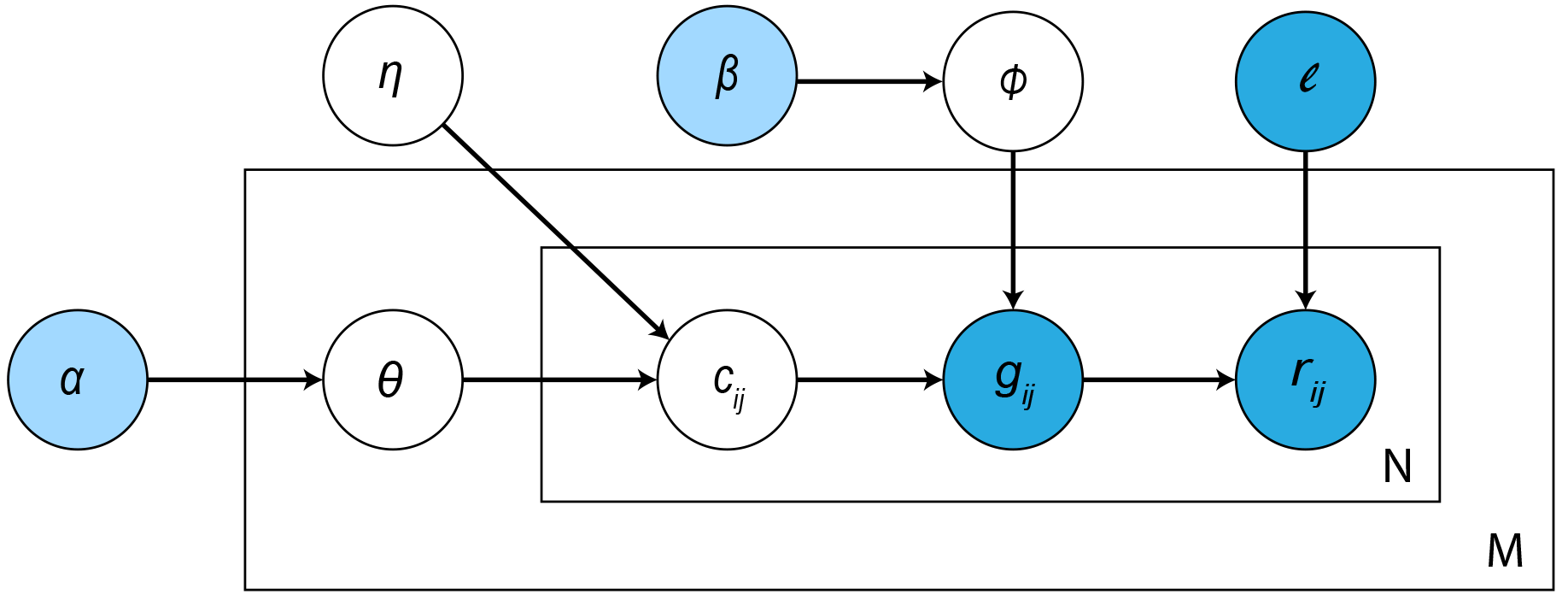
Graphical representation of CDSeq probabilistic model. The cell-type-specific GEPs *ϕ* are modeled by a Dirichlet distribution with hyperparameter *β*, namely, the gene expression profile for cell type *t*, *ϕ*_*t*_ ~ *Dir*(*β*). Our model assumes the vector of sample-specific cell-type mixing proportions *θ* is characterized by another Dirichlet random variable for sample *i*, *ϕ*_*i*_ ~ *Dir*(*α*). A read is then generated as a consequence of the following three consecutive steps: determine the cell type *c*_*i,j*_ from which the read *r*_i,j_ will be produced based on a multinomial distribution with parameter *θ*_*i*_; select a gene *g*_*i,j*_ to which the read *r*_*i,j*_ maps according to a multinomial distribution governed by *ϕ*_*t*_; generate read *r*_*i,j*_ from gene *g*_*i,j*_ with probability determined by the length of gene *ℓ*_*g*_, and Poisson random variables parameters 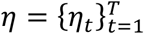(cell size model). The light blue nodes denote the hyperparameters that are assumed to be known. The dark blue nodes denote the values of observable random variables (either measured in the study or established in previous studies) whereas the white nodes are unobservable random variables that need to be inferred from data. The outer box represents samples where M is the sample size, and the inner box denotes the RNA-Seq data of a sample where N is the total number of reads from the sample.

### Statistical inference

Based on our model, given the hyperparameters, reads alignment, effective length of the mapped genes and the estimated value of *η*, the joint distribution of the parameters of interest is given in the following,

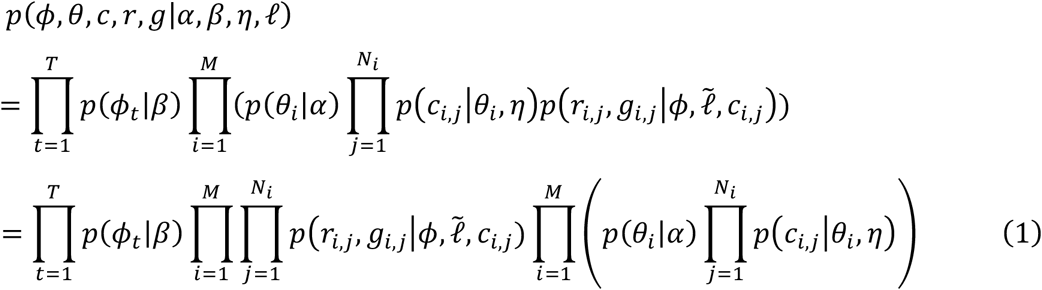

Integrating out *θ* and *ϕ*, we have

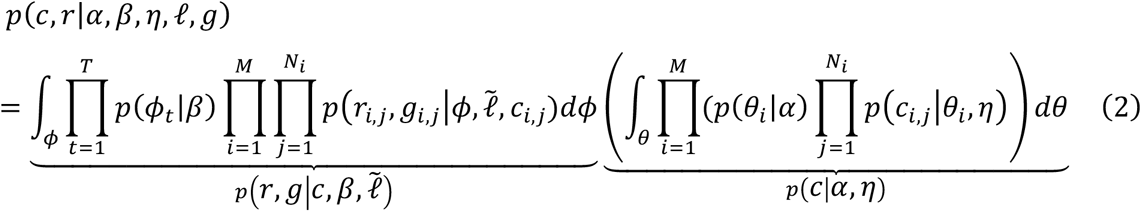

Our goal is to evaluate 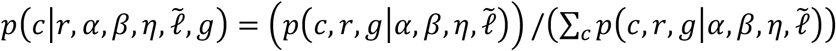 however, we can only evaluate this quantity up to a normalizing constant, i.e., 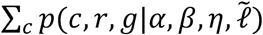. Therefore, we employed a Gibbs sampler^30^, a Markov chain Monte Carlo (MCMC) method, to draw samples from 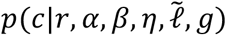, the posterior distribution of cell type assignments, and then use the cell-type assignment information to estimate *θ* and *ϕ*. The Gibbs sampler runs on the space of all possible cell-type assignments for all reads from all the samples. After running MCMC iterations, one can take the value from last MCMC iteration as the cell-type assignment for each read for estimating the model parameters (see Supplementary on statistical inference for details).

#### Determine the number of cell types using the data

The number of cell types present in a heterogenous sample may not be known in advance. Our model allows the number of cell types to be inferred from the data. We formulate this inference as a problem of model selection^30,31^. Based on our statistical model, assuming that the hyperparameters *α*, *β* and the reads mapping information are known, the joint probability distribution of the cell-type-specific GEPs, the sample-specific cell-type proportions in all samples, and the mapped reads is given in eqn.(3) (see Supplementary on statistical inference for details):

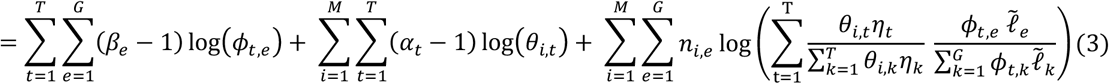

Then, we run CDSeq for a sequence of different numbers of cell types (b, in eqn. 3) and choose the one that maximizes log posterior of eqn.(3). Notice that the minimum number of cell types is two instead of one since one cell type indicates a trivial case of cell type assignment of RNA-seq reads and domain of posterior 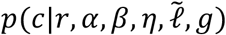 will shrink to a single point.

#### Cell type association

The output of CDSeq reports cell types that are anonymous -- in the sense of not being identified with actual cell types. To match the CDSeq-identified cell types to actual cell type, a list of reference cell-type-specific GEPs and metric of similarity (for example, Pearson’s correlation coefficient or Kullback-Leibler divergence) is required. We employed Pearson’s correlation coefficient as the similarity measurement. Ideally, each estimated cell type will have a high correlation (say exceeding 0.6) with exactly one reference cell type so that the matching is straightforward. In practice, the CDSeq-identified cell types cannot always be uniquely assigned to actual cell types. In such cases, the Munkres algorithm^32^ can be employed to yield one-to-one cell type associations. In some cases, one-to-one cell type association is not immediate (**Figure 6**, **Figure 7**), because a CDSeq-identified cell type may highly correlated with multiple actual cell types. Then, the sample-specific proportion for an actual cell type can be estimated by combining all the proportions of the CDSeq-identified cell types that are highly correlated (say greater than 0.6 or other user defined threshold) with that cell type. We adopted root mean square error (RMSE) as the performance assessment measurement which is defined as

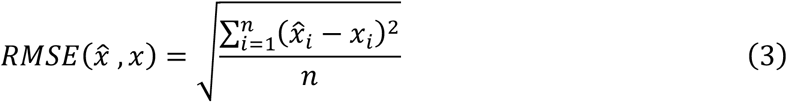

where 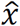 denotes the estimated parameter, *x* denotes the true parameter and *n* denotes the dimension of the parameter. In all the comparisons, we computed the RMSE of estimated GEPs in the original RPKM scale instead of log2 scale.

### A quasi-supervised learning strategy

CDSeq is an unsupervised learning method that aims at discovering the latent pattern from data without any labeling or prior knowledge. The advantage of unsupervised learning framework is that it may unveil some novel information, whereas a limitation is that, when data are too complex and involve sources of variation that are not considered in the model, solution space may be multi-modal. In such cases, CDSeq may find only local optima and the solutions may be hard to interpret biologically. Such scenarios have less impact in a field like text mining than they do in biological research. In text mining, the topics that LDA identifies are abstract notions without any corresponding ground truth. The major task for LDA is to extract latent information that can describe the documents -- there is no basis for deciding whether a topic identified by LDA is actually a “true” topic or not. Instead, in biology, cell types do exist as entities in the mixtures and represent a ground truth that we want to discover. As we showed in our examples, the cell types identified by CDSeq often correspond unambiguously to pure cell lines with matches derived by high correlation between the two GEPs. Sometimes, however, the GEPs of the CDSeq-identified cell types are not highly correlated with any GEPs of the pure cell line GEPs. This issue may arise because multicollinearity among the GEPs of multiple pure cells complicates the deconvolution problem and renders CDSeq less able to definitively separate cell types. To mitigate this kind of problem, we developed a quasi-unsupervised learning strategy. The idea is to provide CDSeq some guidance that leads the algorithm to more biologically meaningful latent information. The guidance consists of appending a set GEPs of pure cell lines to the original input GEPs of heterogenous samples. The choice of GEPs appended should reflect pure cell lines that are believed to constitute the samples. We showed that, using this quasi-unsupervised strategy, CDSeq provided more informative estimates than those obtained using the fully unsupervised mode (**Figure 6**, **Figure 7**, **Supplementary Figure 6** and **Supplementary Figure 7**). We call this learning strategy quasi-unsupervised because, although we do not incorporate any labeling information within CDSeq algorithm itself, we do inject strong signals about likely relevant cell types into the input data. In short, CDSeq is not explicitly aware of such labeling information (pure cell line GEPs appended to input) unlike traditional semi-supervised methods where the labeling information is explicitly taken into account by the algorithms.

### A data dilution strategy to speed up the algorithm

Often data sets of RNA-seq raw counts are large; the total reads across all samples could range from millions to billions. Since CDSeq’s Gibbs sampler is running on the space of all the raw reads, excessively large data sets could dramatically slow the algorithm or even kill it if memory requirements exceed capacity. To address this issue, we propose a data dilution strategy: we divide all the read counts by a positive constant and round them to integer values (since CDSeq requires positive integers as input). We showed that our dilution strategy can speed up the algorithm while retaining the accuracy of estimation. Specifically, we tested this dilution strategy on the synthetic data set by dividing read counts by a sequence of positive integers (100 to 5000 with increment 50). We started from 100 instead of 1 because the original data, which requires around 70GB memory, exceeded the capacity of our hardware and because we wanted to start with a data set that could run in hours instead of days. Dividing by 100 yielded data that took less than 3 hours to run and provided accurate results. We recorded the running time and the correlations between estimated GEP and true GEP (**Supplementary Figure 8**). As we observed, the running time dropped dramatically as the divisor increased from 100 to 500 while the correlations remain relatively close to 1. Taking advantage of this strategy, we could speed up the running time with little compromise in accuracy.

### Deconvolution methods for comparison

We compared our results to those of csSAM^1^ and CIBERSORT^23^, two state-of-the-art computational deconvolution methods. csSAM assumes that expression value of a gene in bulk tissue is the weighted sum of the gene expression in each component within the bulk tissue plus a random error, then given the proportions, i.e., the weight for each component, csSAM fits the linear model by a standard least-square regression of the bulk tissue expression levels on the given proportions to yield the estimated gene expression of each component. We downloaded the csSAM R package and used the default settings for all simulations and comparisons. On the other hand, CIBERSORT, based on the same linearity argument, applies a support vector regression method, called *ν*–support vector regression^33^. It takes as input the gene expression profile of the bulk tissue samples and a gene expression profile for each possible cell type that comprises the bulk tissue; it outputs an estimate of the cell-type proportions for each sample. In addition, to study the fractions of immune cells, a gene expression signature profile for 22 cell types, named LM22, was proposed by CIBERSORT. We requested the source code for CIBERSORT from the authors and ran all comparisons with default settings. Note that CDSeq is not the first method that related to LDA, the methods proposed by ^19,20,27^ are also based on LDA, however, their methods require pure cell line GEPs as input and allow the cell-type-specific GEPs to vary across samples. Therefore, those methods essentially perform partial deconvolution instead of complete deconvolution.

## Supporting information

## Acknowledgments

We are grateful to Dr. Jiajia Wang and Dr. Zongli Xu for their comments and suggestions. We thank Dr. Jiajia Wang for the helpful discussions on the choice of software for processing raw fastq data. We thank the Integrative Bioinformatics Support Group and the Epigenomics Core for the assistance on RNA sequencing and data quality control. We thank the office of the Scientific Information Officer (SIO) at NIEHS and the Computational Biology Facility for computing time. This research was supported in part by Intramural Research Program of the National Institutes of Health, National Institute of Environmental Health Sciences (ES101765).

## Author Contributions

K.K. and L.L. conceived of CDSeq. K.K. developed the model and solver, implemented the algorithm, and generated the synthetic data. L.L., I.S., D.U., K.K. X.L. and Y.L. designed the wet lab experiment. I.S., and M.L. performed the wet lab experiments. K.K. and Q.M. performed data analysis. K.K., D.U., I.S., L.L, Y.L., X.L. and Q.M. wrote the manuscript.

## Supplementary Figures and Tables

**Supplementary Table 1.**
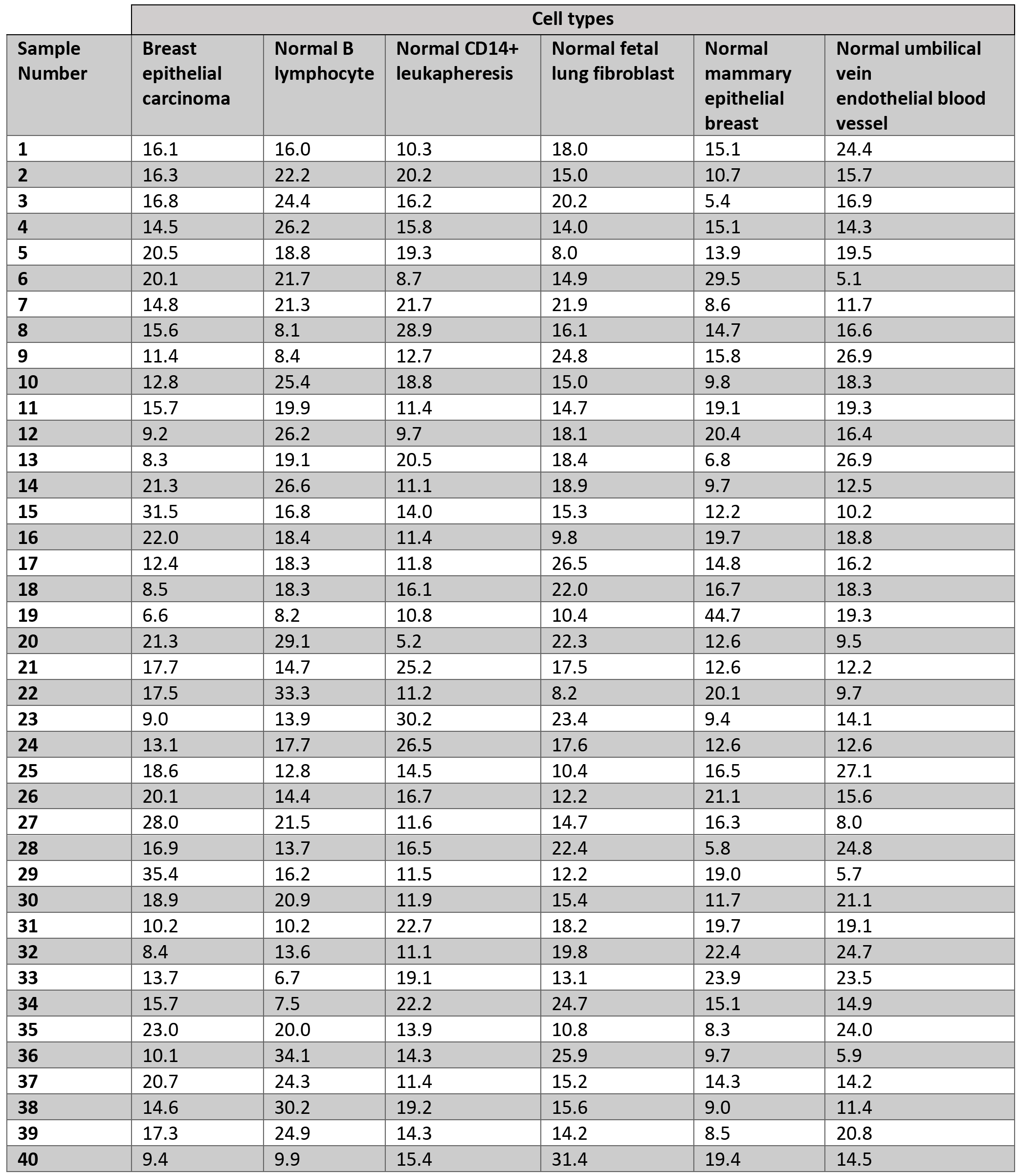
Randomly generated sample-specific cell-type proportions (%) used to create synthetic data.

| Sample Number | Cell types |  |  |  |  |  |
| --- | --- | --- | --- | --- | --- | --- |
|  | Breast epithelial carcinoma | Normal B lymphocyte | Normal CD14+ leukapheresis | Normal fetal lung fibroblast | Normal mammary epithelial breast | Normal umbilical vein endothelial blood vessel |
| 1 | 16.1 | 16.0 | 10.3 | 18.0 | 15.1 | 24.4 |
| 2 | 16.3 | 22.2 | 20.2 | 15.0 | 10.7 | 15.7 |
| 3 | 16.8 | 24.4 | 16.2 | 20.2 | 5.4 | 16.9 |
| 4 | 14.5 | 26.2 | 15.8 | 14.0 | 15.1 | 14.3 |
| 5 | 20.5 | 18.8 | 19.3 | 8.0 | 13.9 | 19.5 |
| 6 | 20.1 | 21.7 | 8.7 | 14.9 | 29.5 | 5.1 |
| 7 | 14.8 | 21.3 | 21.7 | 21.9 | 8.6 | 11.7 |
| 8 | 15.6 | 8.1 | 28.9 | 16.1 | 14.7 | 16.6 |
| 9 | 11.4 | 8.4 | 12.7 | 24.8 | 15.8 | 26.9 |
| 10 | 12.8 | 25.4 | 18.8 | 15.0 | 9.8 | 18.3 |
| 11 | 15.7 | 19.9 | 11.4 | 14.7 | 19.1 | 19.3 |
| 12 | 9.2 | 26.2 | 9.7 | 18.1 | 20.4 | 16.4 |
| 13 | 8.3 | 19.1 | 20.5 | 18.4 | 6.8 | 26.9 |
| 14 | 21.3 | 26.6 | 11.1 | 18.9 | 9.7 | 12.5 |
| 15 | 31.5 | 16.8 | 14.0 | 15.3 | 12.2 | 10.2 |
| 16 | 22.0 | 18.4 | 11.4 | 9.8 | 19.7 | 18.8 |
| 17 | 12.4 | 18.3 | 11.8 | 26.5 | 14.8 | 16.2 |
| 18 | 8.5 | 18.3 | 16.1 | 22.0 | 16.7 | 18.3 |
| 19 | 6.6 | 8.2 | 10.8 | 10.4 | 44.7 | 19.3 |
| 20 | 21.3 | 29.1 | 5.2 | 22.3 | 12.6 | 9.5 |
| 21 | 17.7 | 14.7 | 25.2 | 17.5 | 12.6 | 12.2 |
| 22 | 17.5 | 33.3 | 11.2 | 8.2 | 20.1 | 9.7 |
| 23 | 9.0 | 13.9 | 30.2 | 23.4 | 9.4 | 14.1 |
| 24 | 13.1 | 17.7 | 26.5 | 17.6 | 12.6 | 12.6 |
| 25 | 18.6 | 12.8 | 14.5 | 10.4 | 16.5 | 27.1 |
| 26 | 20.1 | 14.4 | 16.7 | 12.2 | 21.1 | 15.6 |
| 27 | 28.0 | 21.5 | 11.6 | 14.7 | 16.3 | 8.0 |
| 28 | 16.9 | 13.7 | 16.5 | 22.4 | 5.8 | 24.8 |
| 29 | 35.4 | 16.2 | 11.5 | 12.2 | 19.0 | 5.7 |
| 30 | 18.9 | 20.9 | 11.9 | 15.4 | 11.7 | 21.1 |
| 31 | 10.2 | 10.2 | 22.7 | 18.2 | 19.7 | 19.1 |
| 32 | 8.4 | 13.6 | 11.1 | 19.8 | 22.4 | 24.7 |
| 33 | 13.7 | 6.7 | 19.1 | 13.1 | 23.9 | 23.5 |
| 34 | 15.7 | 7.5 | 22.2 | 24.7 | 15.1 | 14.9 |
| 35 | 23.0 | 20.0 | 13.9 | 10.8 | 8.3 | 24.0 |
| 36 | 10.1 | 34.1 | 14.3 | 25.9 | 9.7 | 5.9 |
| 37 | 20.7 | 24.3 | 11.4 | 15.2 | 14.3 | 14.2 |
| 38 | 14.6 | 30.2 | 19.2 | 15.6 | 9.0 | 11.4 |
| 39 | 17.3 | 24.9 | 14.3 | 14.2 | 8.5 | 20.8 |
| 40 | 9.4 | 9.9 | 15.4 | 31.4 | 19.4 | 14.5 |

**Supplementary Table 2.** Cell-type (RNA) proportions (%) used to create mixed samples in the experiment with cultured cell types. Samples 1 to 4 and 37 to 40 are two replicates of the four pure cell lines. They were used as ground truth for benchmarking the CDSeq-identified cell types, not input for CDSeq.

| Sample Number | Cell types |  |  |  |
| --- | --- | --- | --- | --- |
|  | Tumor – MCF7 | CAFs – Hs 343.T | Normal breast – hMECs-hTERT | Lymphocytes – Namalwa |
| 1 | 100 | 0 | 0 | 0 |
| 2 | 0 | 100 | 0 | 0 |
| 3 | 0 | 0 | 100 | 0 |
| 4 | 0 | 0 | 0 | 100 |
| 5 | 85 | 5 | 5 | 5 |
| 6 | 85 | 9 | 3 | 3 |
| 7 | 85 | 3 | 9 | 3 |
| 8 | 85 | 3 | 3 | 9 |
| 9 | 70 | 10 | 10 | 10 |
| 10 | 70 | 15 | 10 | 5 |
| 11 | 70 | 15 | 5 | 10 |
| 12 | 70 | 10 | 15 | 5 |
| 13 | 70 | 10 | 5 | 15 |
| 14 | 70 | 5 | 15 | 10 |
| 15 | 70 | 5 | 10 | 15 |
| 16 | 55 | 15 | 15 | 15 |
| 17 | 55 | 30 | 10 | 5 |
| 18 | 55 | 30 | 5 | 10 |
| 19 | 55 | 10 | 30 | 5 |
| 20 | 55 | 10 | 5 | 30 |
| 21 | 55 | 5 | 30 | 10 |
| 22 | 55 | 5 | 10 | 30 |
| 23 | 40 | 20 | 20 | 20 |
| 24 | 40 | 30 | 20 | 10 |
| 25 | 40 | 30 | 10 | 20 |
| 26 | 40 | 20 | 30 | 10 |
| 27 | 40 | 20 | 10 | 30 |
| 28 | 40 | 10 | 30 | 20 |
| 29 | 40 | 10 | 20 | 30 |
| 30 | 25 | 25 | 25 | 25 |
| 31 | 25 | 35 | 25 | 15 |
| 32 | 25 | 35 | 15 | 25 |
| 33 | 25 | 25 | 35 | 15 |
| 34 | 25 | 25 | 15 | 35 |
| 35 | 25 | 15 | 35 | 25 |
| 36 | 25 | 15 | 25 | 35 |
| 37 | 100 | 0 | 0 | 0 |
| 38 | 0 | 100 | 0 | 0 |
| 39 | 0 | 0 | 100 | 0 |
| 40 | 0 | 0 | 0 | 100 |

**Supplementary Figure 1.**
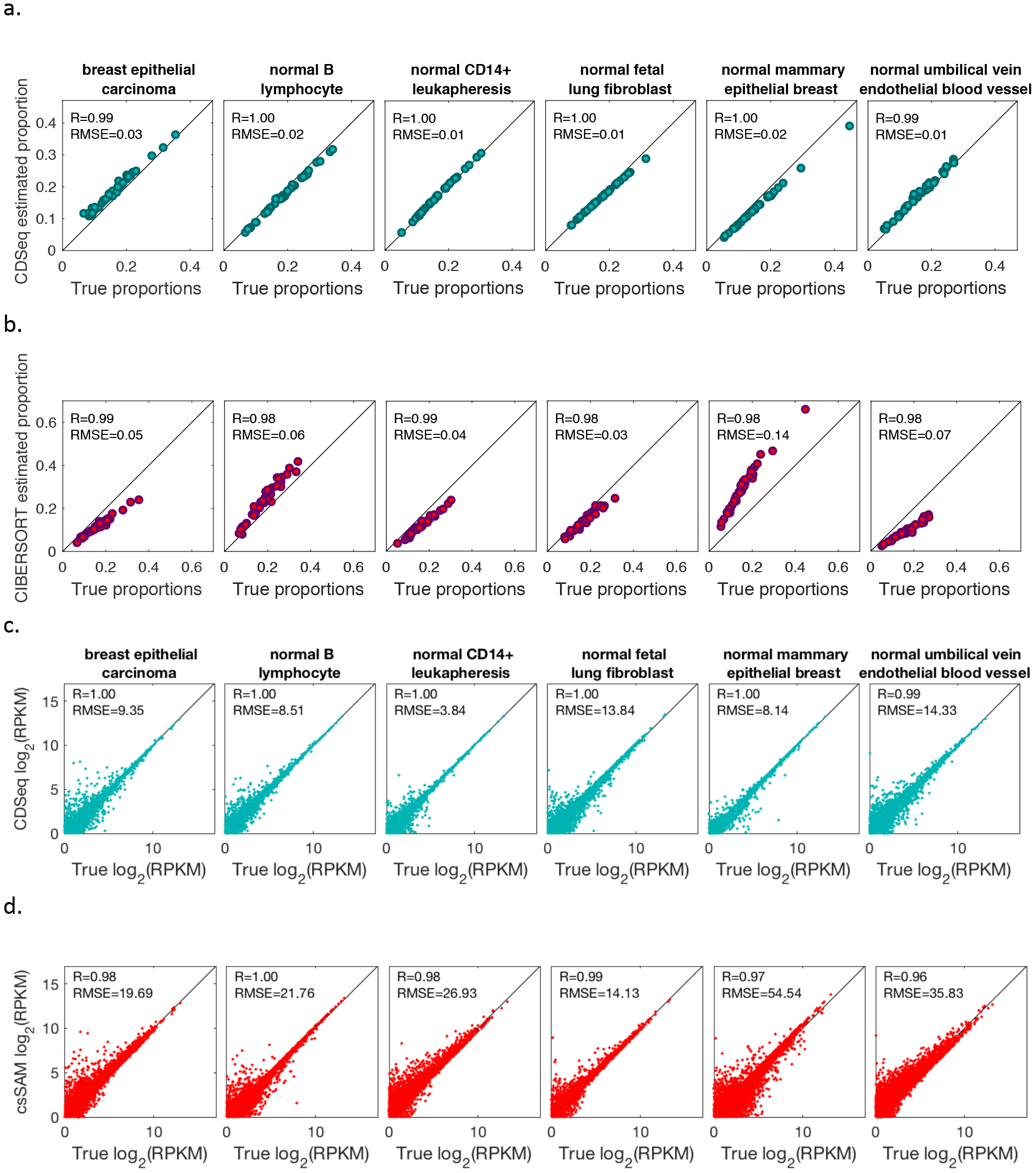
Results for synthetic data. (a). CDSeq estimates versus true proportions; (b). CIBERSORT estimates versus true proportions; (c).CDSeq GEP estimates versus true GEP; (d). csSAM GEP estimates versus true GEP.

**Supplementary Figure 2.**
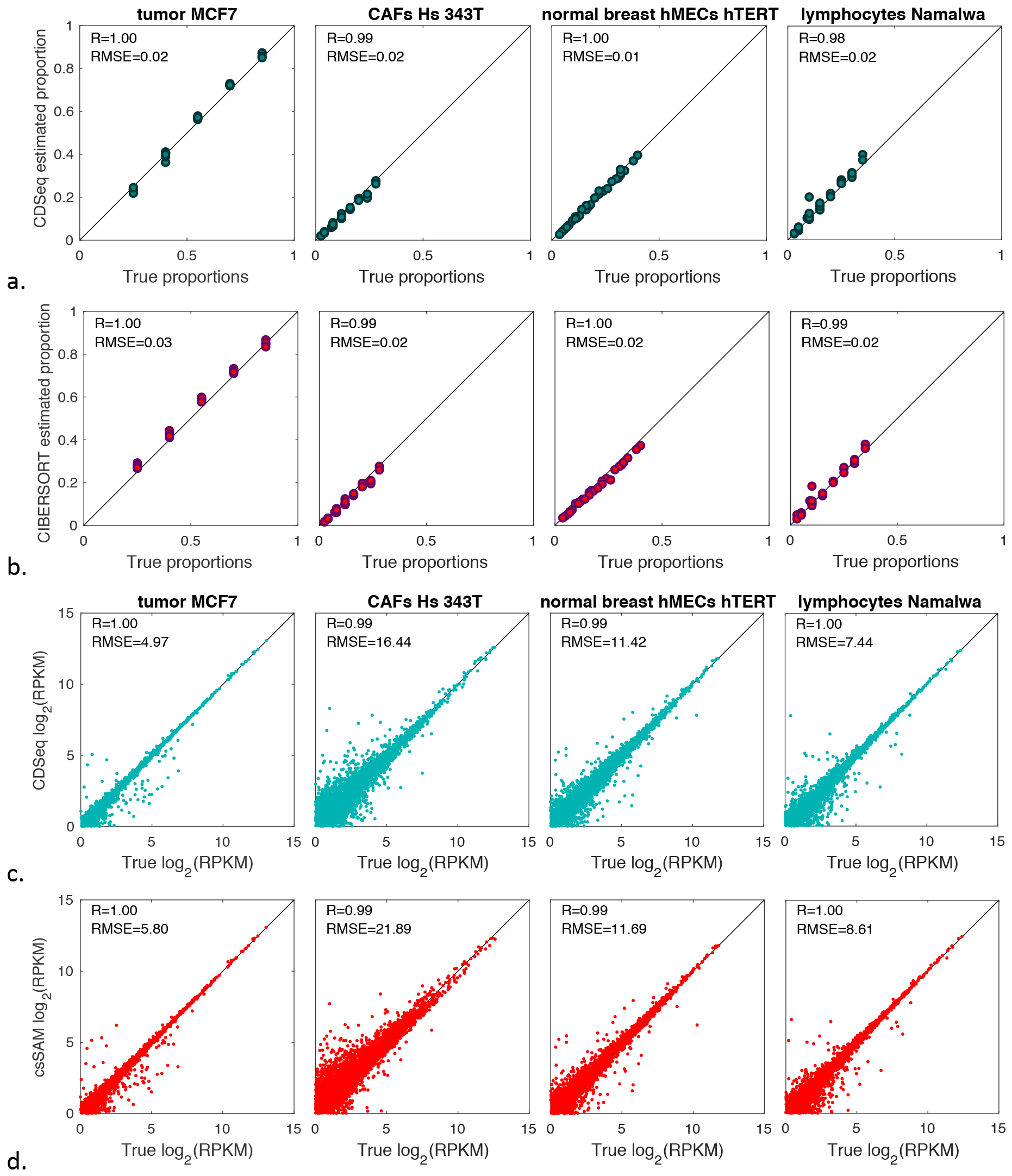
Results from deconvolution of experimental data: mixed RNA from cultured cell lines. (a). CDSeq estimated proportions versus true proportions; (b). CIBERSORT estimated proportions versus true proportions; (c). CDSeq estimated GEP versus true GEP; (d). csSAM estimated GEP versus true GEP.

**Supplementary Figure 3.**
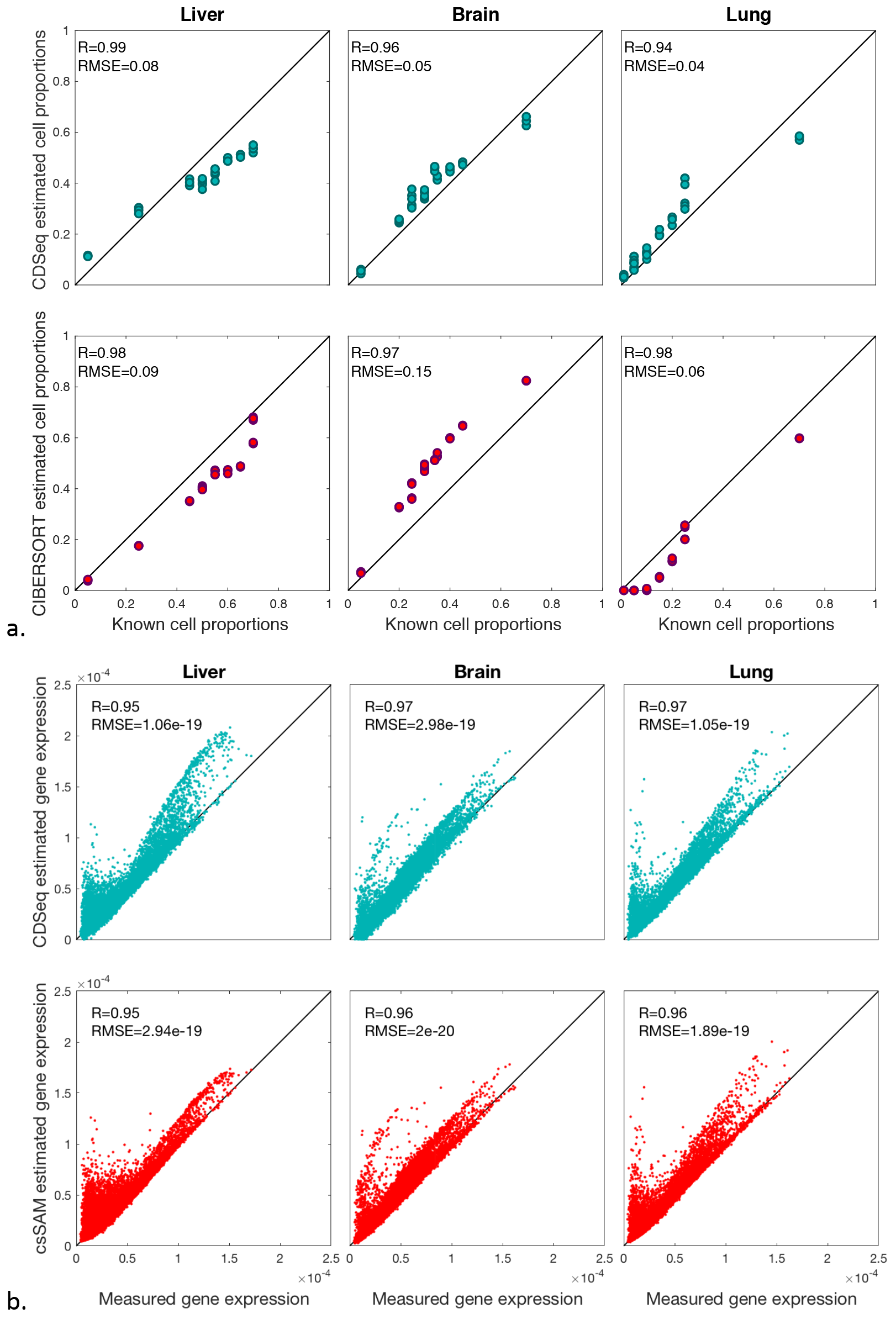
Results from deconvolution of mixed samples of liver, brain and lung cells. (a) Estimated sample-specific cell-type proportions by CDSeq and CIBERSORT compared to known cell-type proportions; (b) Cell-type-specific estimated GEPs by CDSeq and csSAM compared to observed GEPs from pure cell lines.

**Supplementary Figure 4.**
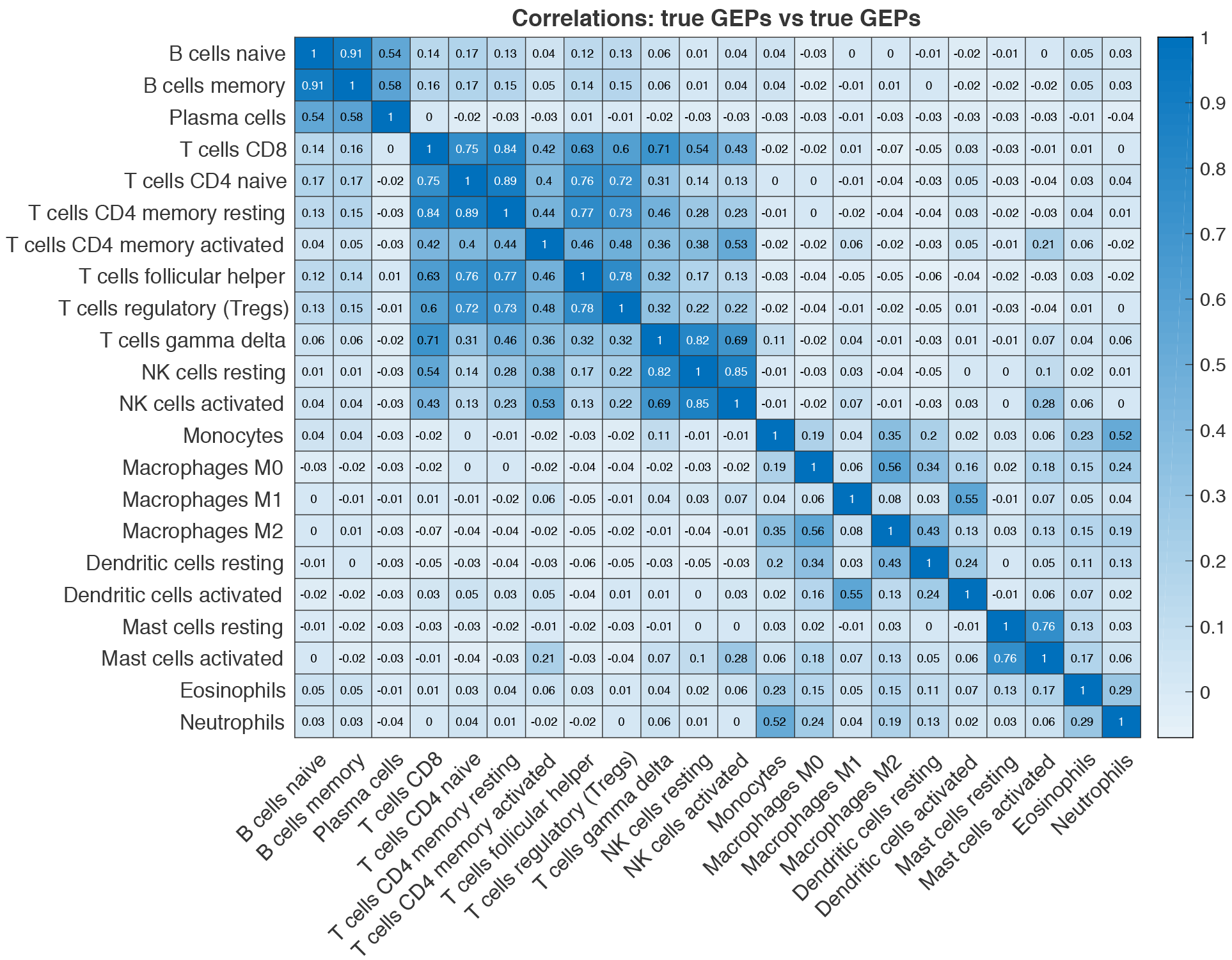
Correlation among true LM22 GEPs. Some of the 22 cell types are highly correlated.

**Supplementary Figure 5.**
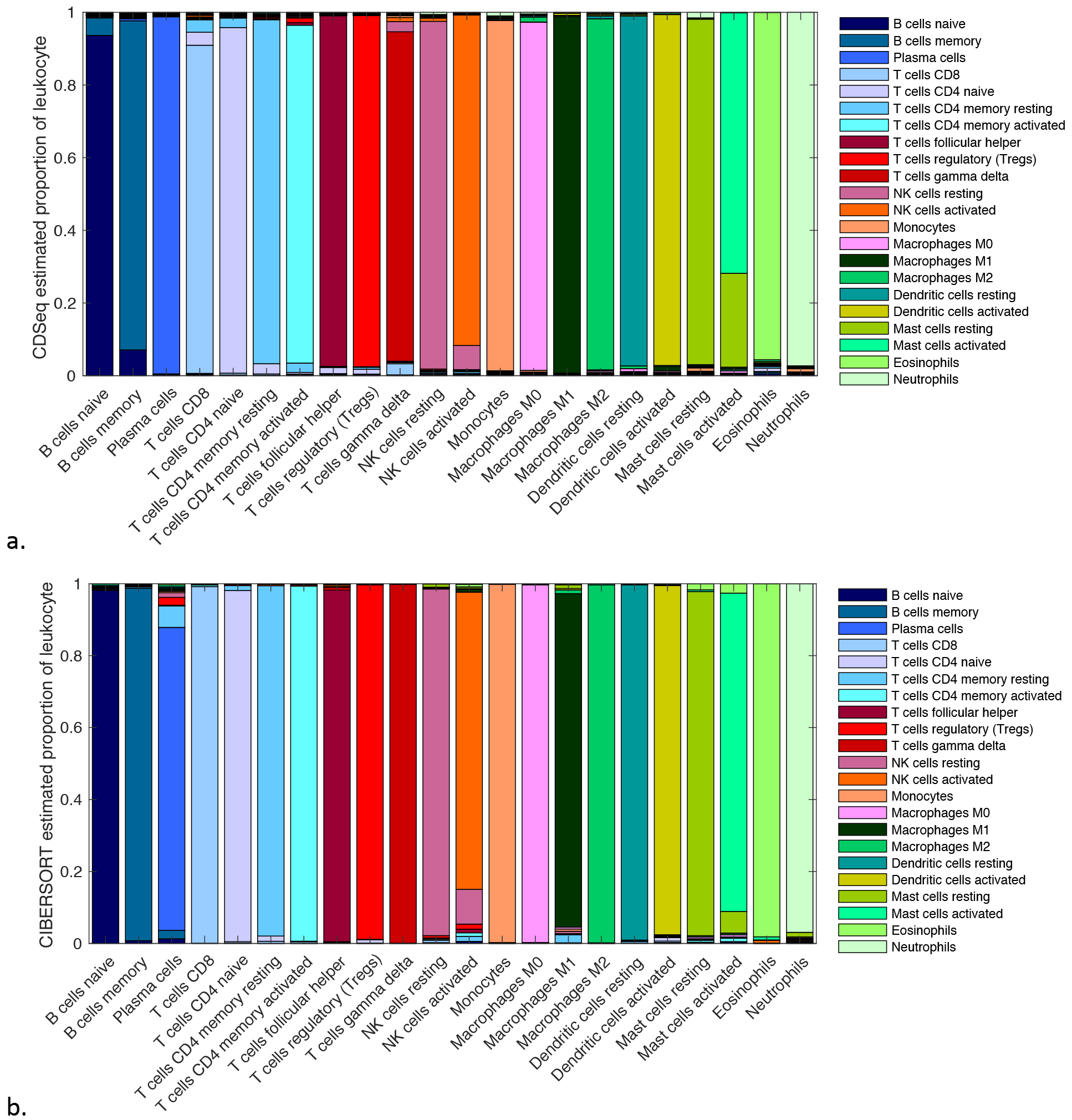
Bar plot of estimated proportions of LM22 data set. Ideally, each bar plot should only contain only a single color since the LM22 is a set of GEPs from pure cell types. The bar plots of both methods are largely, but not exclusively, dominated by single colors, indicating the accuracy of estimated proportions. (a) CDSeq-estimated proportions; (b) CIBERSORT-estimated proportions.

**Supplementary Figure 6.**
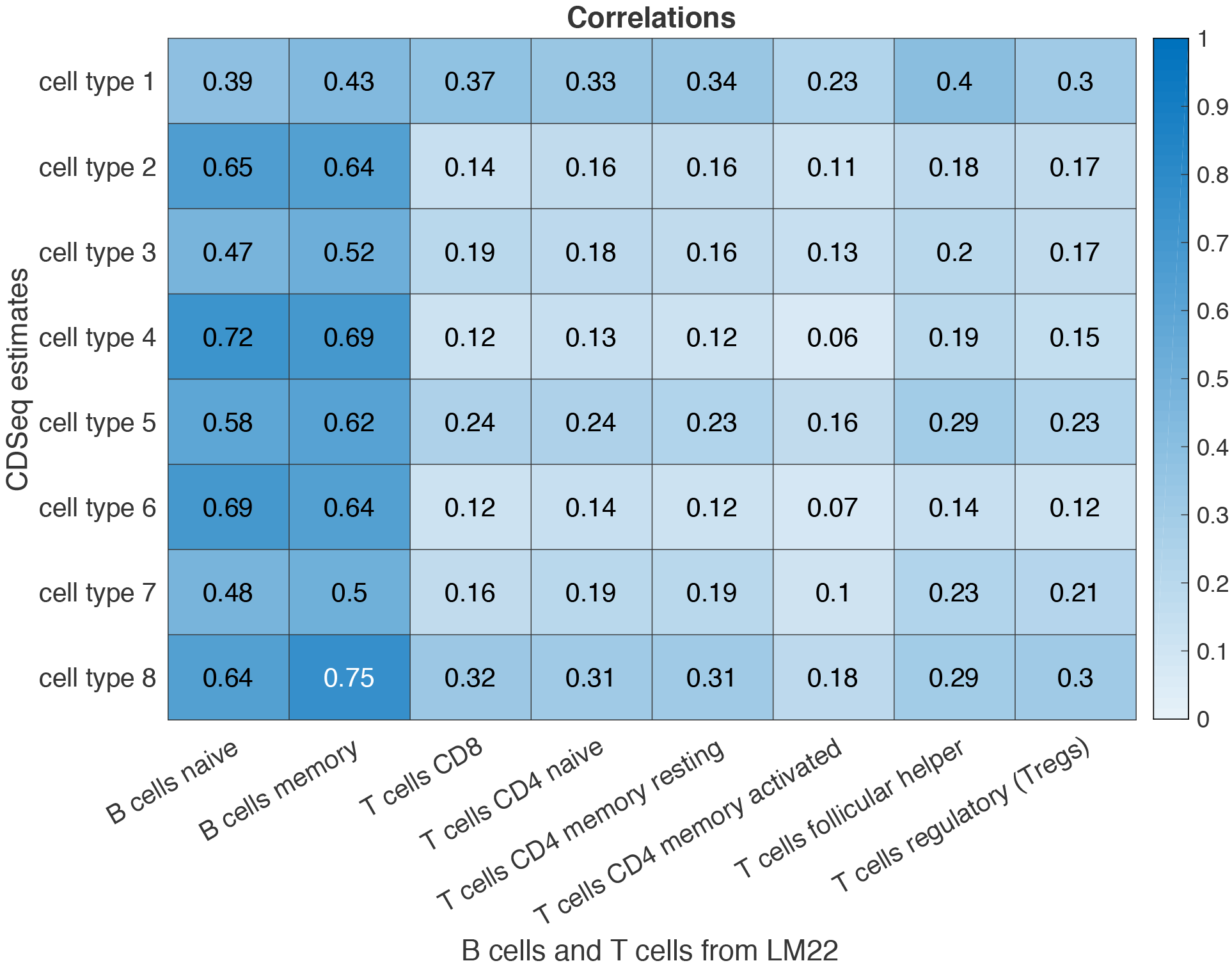
Result of deconvolution of follicular lymphoma tumors data. Heatmap chart of correlations between CDSeq-estimated cell-type-specific GEPs using fully unsupervised mode. We set the number of cell types to be 8, hyperparameters *α* = 5, *β* = 0.5, and 700 MCMC runs. The 8 CDSeq-identified cell-type-specific GEPs were relatively highly correlated with naïve B cell and memory B cell GEPs but less correlated with the T cell GEPs. In fully unsupervised mode, CDSeq could uncover signals only from B cells.

**Supplementary Figure 7.**
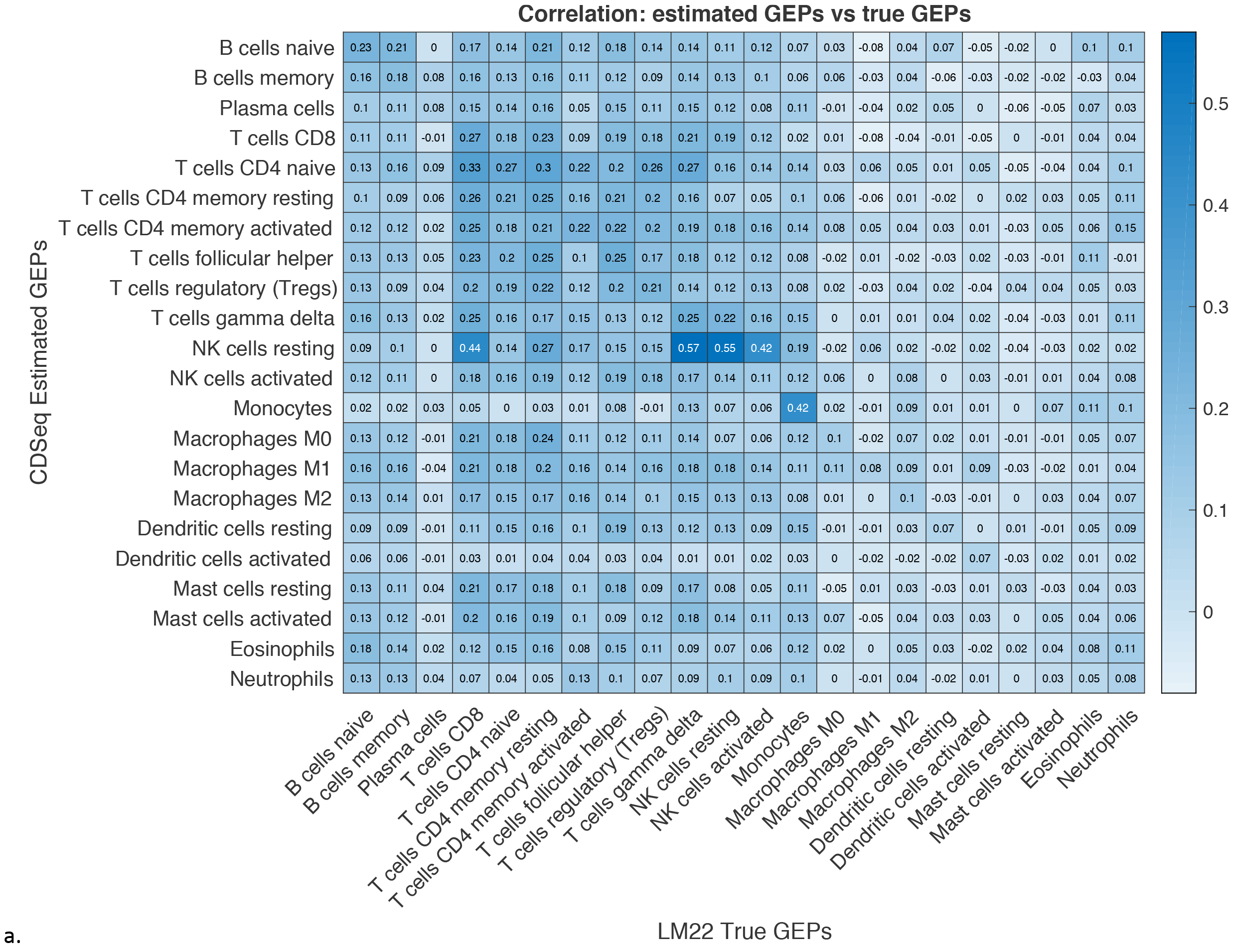
CDSeq performance on deep deconvolution with fully unsupervised learning. We applied CDSeq on PBMC data set 11 without appending LM22 GEPs to the GEPs of the heterogenous samples. We set the number of cell types to be 22, α=50,β=20: (a) Heat map of correlations between estimated cell-type-specific GEPs and LM22 GEPs; and, (b) CDSeq-estimated cell-type proportions compared to flow cytometry estimates, where the upper panel (blue dots) is the result of fully unsupervised mode and the lower panel (green dots) is the result of quasi-unsupervised mode. The black line is the linear regression line and p is the p-value for testing the hypothesis of no correlation against the alternative hypothesis of a nonzero correlation.

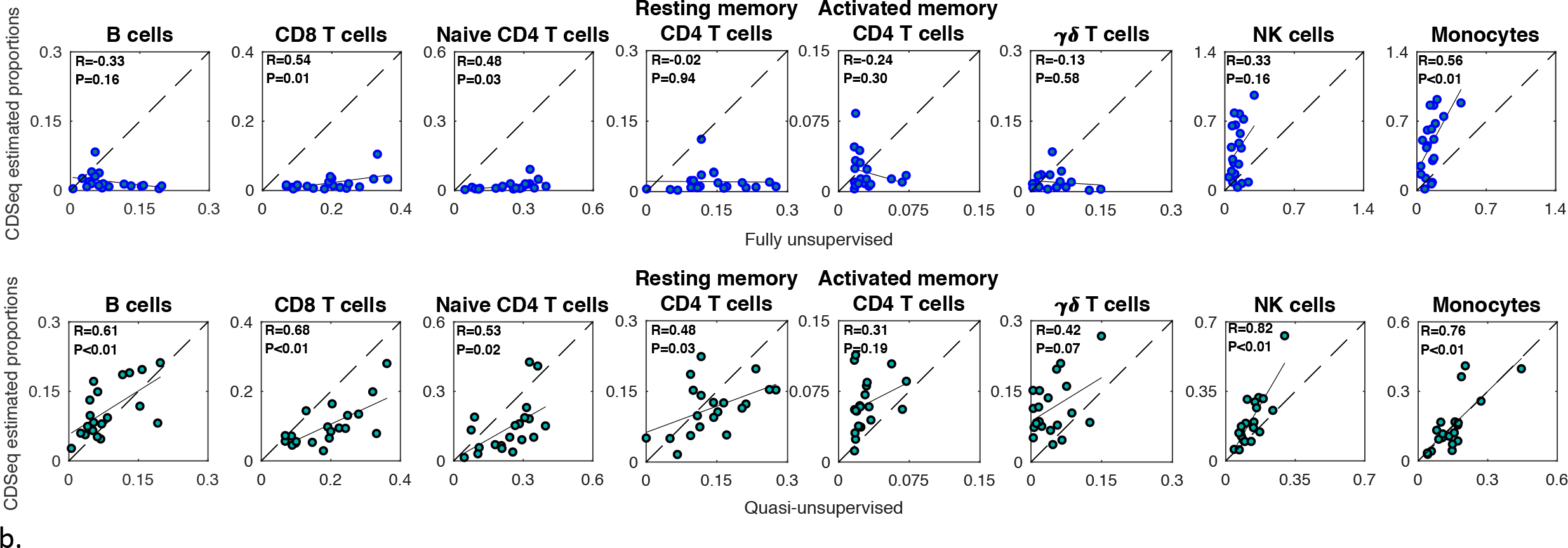

**Supplementary Figure 8.**
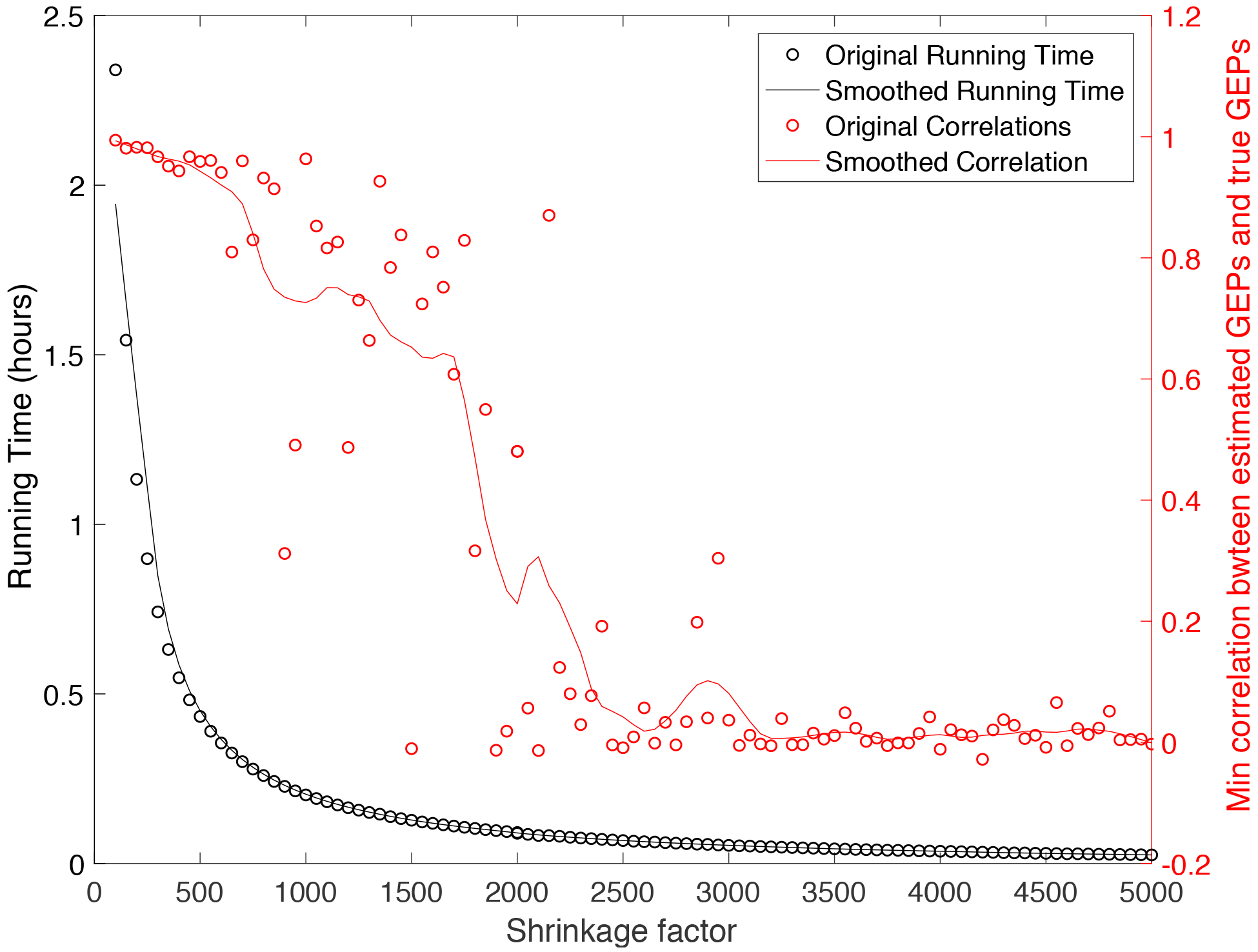
Running time of CDSeq plotted against the shrinkage factor for data dilution using the synthetic data. Data dilution is a strategy to speed up calculations with very large input data sets. One divides the read counts by a constant shrinkage factor and rounds to integer values before running CDSeq. For shrinkage factors ranging from 100 to 5000 in increments of 50, we measured the running time (in hours) and recorded the minimum correlations between the six CDSeq-identified cell-type-specific GEPs and their matched (using Munkres algorithm) six true GEPs. As the shrinkage factor increase, the running time decreases rapidly whereas the minimum of the Pearson correlation coefficients decreases more slowly and reaches a plateau near zero, indicating that at least one of the true cell types could not be identified, after shrinkage factors exceed about 2000. In this example, the total number of reads for all the 40 synthetic samples is about 9.5 × 10^9^. This result shows that large shrinkage factors degrade estimation and impair correct matching of CDSeq-identified cell types to actual cell types.

## Reference

1. Shen-Orr, S. S. et al. Cell type--specific gene expression differences in complex tissues. Nat. Methods 7, 287–289 (2010).

2. Zhong, Y. & Liu, Z. Gene expression deconvolution in linear space. Nat. Methods 9, 8 (2012).

3. Kuhn, A., Thu, D., Waldvogel, H. J., Faull, R. L. M. & Luthi-Carter, R. Population-specific expression analysis (PSEA) reveals molecular changes in diseased brain. Nat. Methods 8, 945–947 (2011).

4. Alizadeh, A. A. et al. Toward understanding and exploiting tumor heterogeneity. Nat. Med. 21, 846–853 (2015).

5. Mlecnik, B. et al. Integrative Analyses of Colorectal Cancer Show Immunoscore Is a Stronger Predictor of Patient Survival Than Microsatellite Instability. Immunity 44, 698–711 (2016).

6. Calon, A. et al. Stromal gene expression defines poor-prognosis subtypes in colorectal cancer. Nat. Genet. (2015). doi:10.1038/ng.3225

7. Galon, J. et al. Towards the introduction of the ‘Immunoscore’ in the classification of malignant tumours. J. Pathol. 232, 199–209 (2014).

8. Zheng, C. et al. Landscape of Infiltrating T Cells in Liver Cancer Revealed by Single-Cell Sequencing. Cell 169, 1342–1356.e16 (2017).

9. Gentles, A. J. et al. The prognostic landscape of genes and infiltrating immune cells across human cancers. Nat. Med. 21, 938–945 (2015).

10. Shen-Orr, S. S. & Gaujoux, R. Computational deconvolution: extracting cell type-specific information from heterogeneous samples. Curr. Opin. Immunol. 25, 571–578 (2013).

11. Hackl, H., Charoentong, P., Finotello, F. & Trajanoski, Z. Computational genomics tools for dissecting tumour-immune cell interactions. Nature Reviews Genetics 17, 441–458 (2016).

12. Gawad, C., Koh, W. & Quake, S. R. Single-cell genome sequencing: Current state of the science. Nature Reviews Genetics 17, 175–188 (2016).

13. Vallejos, C. A., Risso, D., Scialdone, A., Dudoit, S. & Marioni, J. C. Normalizing single-cell RNA sequencing data: Challenges and opportunities. Nature Methods (2017). doi:10.1038/nmeth.4292

14. Avila Cobos, F., Vandesompele, J., Mestdagh, P. & De Preter, K. Computational deconvolution of transcriptomics data from mixed cell populations. Bioinformatics (2018). doi:10.1093/bioinformatics/bty019

15. Venet, D., Pecasse, F., Maenhaut, C. & Bersini, H. Separation of samples into their constituents using gene expression data. Bioinformatics 17, S279-–S287 (2001).

16. Lu, P., Nakorchevskiy, A. & Marcotte, E. M. Expression deconvolution: a reinterpretation of DNA microarray data reveals dynamic changes in cell populations. Proc. Natl. Acad. Sci. 100, 10370–10375 (2003).

17. Erkkilä, T. et al. Probabilistic analysis of gene expression measurements from heterogeneous tissues. Bioinformatics 26, 2571–2577 (2010).

18. Gaujoux, R. & Seoighe, C. Semi-supervised Nonnegative Matrix Factorization for gene expression deconvolution: a case study. Infect. Genet. Evol. 12, 913–921 (2012).

19. Quon, G. & Morris, Q. ISOLATE: a computational strategy for identifying the primary origin of cancers using high-throughput sequencing. Bioinformatics 25, 2882–2889 (2009).

20. Qiao, W. et al. PERT: a method for expression deconvolution of human blood samples from varied microenvironmental and developmental conditions. PLoS Comput. Biol. 8, e1002838 (2012).

21. Gong, T. & Szustakowski, J. D. DeconRNASeq: a statistical framework for deconvolution of heterogeneous tissue samples based on mRNA-Seq data. Bioinformatics 29, 1083–1085 (2013).

22. Li, Y. & Xie, X. A mixture model for expression deconvolution from RNA-seq in heterogeneous tissues. BMC Bioinformatics 14, S11 (2013).

23. Newman, A. M. et al. Robust enumeration of cell subsets from tissue expression profiles. Nat. Methods 12, 453 (2015).

24. Li, B. et al. Comprehensive analyses of tumor immunity: Implications for cancer immunotherapy. Genome Biol. (2016). doi:10.1186/s13059-016-1028-7

25. Li, B., Liu, J. S. & Liu, X. S. Revisit linear regression-based deconvolution methods for tumor gene expression data. Genome Biol. (2017). doi:10.1186/s13059-017-1256-5

26. Blei, D. M., Ng, A. Y. & Jordan, M. I. Latent dirichlet allocation. J. Mach. Learn. Res. 3, 993–1022 (2003).

27. Quon, G. et al. Computational purification of individual tumor gene expression profiles leads to significant improvements in prognostic prediction. Genome Med. 5, 29 (2013).

28. Marguerat, S. & Bähler, J. Coordinating genome expression with cell size. Trends in Genetics (2012). doi:10.1016/j.tig.2012.07.003

29. Zhong, Y., Wan, Y.-W., Pang, K., Chow, L. M. L. & Liu, Z. Digital sorting of complex tissues for cell type-specific gene expression profiles. BMC Bioinformatics 14, 89 (2013).

30. Griffiths, T. L. & Steyvers, M. Finding scientific topics. Proc. Natl. Acad. Sci. 101, 5228–5235 (2004).

31. Kass, R. R. E. & Raftery, A. E. A. Bayes factors. J. Am. Stat. Assoc.… 90, 773–795 (1995).

32. Burkard, R. E., Dell’Amico, M. & Martello, S. Assignment problems, revised reprint. 125, (Siam, 2009).

33. Schölkopf, B., Smola, A. J., Williamson, R. C. & Bartlett, P. L. New support vector almgorithms. Neural Comput. 12, 1207–1245 (2000).

