## Supplementary material for "A novel computational complete deconvolution method using RNA-seq data"

**We introduce CDSeq, a novel computational method for complete deconvolution of tissue samples, where each sample is a heterogeneous mixture of cell types. Using RNA-seq gene expression profiles from a collection of samples, our method estimates cell-type-specific expression profiles in common across samples as well as sample-specific proportions of each cell type. We benchmarked CDSeq using several synthetic and experimental datasets, and we compared its performance with two state-of-the-art partial deconvolution methods. CDSeq performed as well or better than the other methods in estimating the cell-type-specific gene expression profiles and sample-specific proportions of cell types.**

### 1 Supplementary materials on statistical inference

| Notations | |
|---|---|
| $M$ | Number of tissue samples |
| $G$ | Total number of genes |
| $N_i$ | Total number of reads for sample $i$ |
| $N_t$ | Total number of reads assigned to cell type $t$ for all samples |
| $\theta_i$ | Multinomial parameters that describe the sample-specific proportions of the cell types for sample $i$, $\theta_i \in \mathbb{R}_+^T$, $\sum_{t=1}^{T} \theta_{i,t} = 1$and let $\theta = (\theta_{i,t})_{i=1,t=1}^{M,T}$ |
| $\phi_t$ | Multinomial parameters that describe the cell-type-specific gene expression for cell type $t$, $\phi_t \in \mathbb{R}_+^G$, $\sum_{e=1}^{G} \phi_{t,e} = 1$ and let $\phi = (\phi_{t,e})_{t=1,e=1}^{T,G}$ |
| $\alpha = (\alpha_i)_{i=1}^M$ | Dirichlet parameter, hyperparameter for $\theta_i$ |
| $\beta = (\beta_e)_{e=1}^G$ | Dirichlet parameter, hyperparameter for $\phi_t$ |
| $r_{i,j}$ | mapped RNA-seq read $j$ for sample $i$, let $r = (r_{i,j})_{i=1,j=1}^{M,N_i}$ |
| $c_{i,j}$ | cell type assignment for read $r_{i,j}$, $c_{i,j} \in \{1, \cdots, T\}$ and let $c = (c_{i,j})_{i=1,j=1}^{M,N_i}$ |
| $\eta_t$ | Poisson parameter that describes the amount of RNA generated from cell type $t$, let $\eta = (\eta_t)_{t=1}^T$ |
| $\ell_e$ | gene length for gene $e$, let $\ell = (\ell_e)_{e=1}^G$ |
| $\tilde{\ell}_e$ | effective gene length for gene $e$, $\tilde{\ell}_e = \ell_e - m + 1$, where $m$ is the fixed sequencing length, let $\tilde{\ell} = (\tilde{\ell}_e)_{e=1}^G$ |
| $g_{i,j}$ | mapped gene of read $r_{i,j}$, $g_{i,j} \in \{1, \cdots, G\}$ and let $g = (g_{i,j})_{i=1,j=1}^{M,N_i}$ |

| Notations (continue) | |
|---|---|
| $gid(r_a)$ | mapping from a read $r_a$ to its gene. We sometimes dropped the double subscripts $i, j$ for convenience and use $r = \{r_1, \cdots, r_n\}$ to denote all the reads from all samples |
| $cid(r_a)$ | mapping from a read $r_a$ to its cell type assignment |
| $sid(r_a)$ | mapping from a read $r_a$ to its sample |

For our model (**Methods**), given the hyperparameters, reads alignment, effective lengths of the mapped genes and the estimate of $\eta$, the joint distribution of the parameters of interest is

$$\begin{aligned} & p(\phi, \theta, c, r, g|\alpha, \beta, \eta, \tilde{\ell}) \\ = & \prod_{t=1}^{T} p(\phi_t|\beta) \prod_{i=1}^{M} \left( p(\theta_i|\alpha) \prod_{j=1}^{N_i} p(c_{i,j}|\theta_i, \eta) p(r_{i,j}, g_{i,j}|\phi, \tilde{\ell}, c_{i,j}) \right) \\ = & \prod_{t=1}^{T} p(\phi_t|\beta) \prod_{i=1}^{M} \prod_{j=1}^{N_i} p(r_{i,j}, g_{i,j}|\phi, \tilde{\ell}, c_{i,j}) \prod_{i=1}^{M} \left( p(\theta_i|\alpha) \prod_{j=1}^{N_i} p(c_{i,j}|\theta_i, \eta) \right). \end{aligned}$$

Integrating out $\theta$ and $\phi$, we have

$$\begin{aligned} & p(c, r, g|\alpha, \beta, \eta, \tilde{\ell}) \\ = & \underbrace{\int_{\phi} \prod_{t=1}^{T} p(\phi_t|\beta) \prod_{i=1}^{M} \prod_{j=1}^{N_i} p(r_{i,j}, g_{i,j}|\phi, \tilde{\ell}, c_{i,j}) d\phi}_{p(r,g|c,\beta,\tilde{\ell})} \underbrace{\int_{\theta} \prod_{i=1}^{M} \left( p(\theta_i|\alpha) \prod_{j=1}^{N_i} p(c_{i,j}|\theta_i, \eta) \right) d\theta}_{p(c|\alpha,\eta)}. \end{aligned}$$

Subsequently,

$$\begin{aligned} & p(r, g|c, \beta, \tilde{\ell}) \\ = & \int_{\phi} \prod_{t=1}^{T} p(\phi_t|\beta) \prod_{i=1}^{M} \prod_{j=1}^{N_i} p(r_{i,j}, g_{i,j}|\phi, \tilde{\ell}, c_{i,j}) d\phi \end{aligned} \quad (1)$$

$$= \prod_{t=1}^{T} \int_{\phi_t} \frac{\Gamma(\sum_{e=1}^{G} \beta_e)}{\prod_{e=1}^{G} \Gamma(\beta_e)} \left( \prod_{e=1}^{G} \phi_{t,e}^{\beta_e - 1} \right) \left( \prod_{\tau=1}^{N_t} \frac{\phi_{t,g_{i_\tau,j_\tau}} \tilde{\ell}_{g_{i_\tau,j_\tau}}}{\sum_{e=1}^{G} \phi_{t,e} \tilde{\ell}_e} \frac{1}{\tilde{\ell}_{g_{i_\tau,j_\tau}}} \right) d\phi_t, \qquad (2)$$

where $N_t$ denotes the number of reads assigned to cell type $t$, and $i_\tau, j_\tau$ denote the sample id and read index within the sample $i_\tau$ respectively, $g_{i_\tau,j_\tau}$ and $\tilde{\ell}_{g_{i_\tau,j_\tau}}$ denote the gene assignment and its effective length from which the read $r_{i_\tau,j_\tau}$ is generated. From (1) to (2), it is done by grouping the reads according to their cell type assignments and writing the Dirichlet distribution for $\phi_t$ explicitly. Similarly, we have that

$$\begin{aligned} & p(c|\alpha, \eta) \\ = & \int_{\theta} \prod_{i=1}^{M} \left( p(\theta_i|\alpha) \prod_{j=1}^{N_i} p(c_{i,j}|\theta_i, \eta) \right) d\theta \\ = & \prod_{i=1}^{M} \int_{\theta_i} \frac{\Gamma(\sum_{t=1}^{T} \alpha_t)}{\prod_{t=1}^{T} \Gamma(\alpha_t)} \left( \prod_{t=1}^{T} \theta_{i,t}^{\alpha_t - 1} \right) \prod_{j=1}^{N_i} \prod_{t=1}^{T} \left( \frac{\theta_{i,t}\eta_t}{\sum_{t=1}^{T} \theta_{i,t}\eta_t} \right)^{\mathbb{1}\{c_{i,j}=t\}} d\theta_i, \qquad (3) \end{aligned}$$

where $\mathbb{1}\{\cdot\}$ is the indicator function takes values of $1$ or $0$. Ideally, one could integrate out the parameters $\theta$ and $\phi$ separately. Nevertheless, such direct computation is complicated by the fact that $p(r_{i,j}, g_{i,j}|\phi, \tilde{\ell}, c_{i,j})$ and $p(c_{i,j}|\theta_i, \eta)$ contain normalizing denominators as shown in (2) and (3).

To simplify the computation, we define $\hat{\phi}_{t,g_{i_\tau,j_\tau}} = \frac{\phi_{t,g_{i_\tau,j_\tau}} \tilde{\ell}_{g_{i_\tau,j_\tau}}}{\sum_{k=1}^{G} \phi_{t,k} \tilde{\ell}_k}$ and $\hat{\theta}_{i,t} = \frac{\theta_{i,t}\eta_t}{\sum_{t=1}^{T} \theta_{i,t}\eta_t}$. Notice that $\hat{\phi}_t$ and $\hat{\theta}_i$ are random variables on simplex. We therefore use two Dirichlet random variables $\tilde{\phi}_t = (\tilde{\phi}_{t,1}, \cdots, \tilde{\phi}_{t,G})$ and $\tilde{\theta}_i = (\tilde{\theta}_{i,1}, \cdots, \tilde{\theta}_{i,T})$, to approximate $\hat{\phi}_t$ and $\hat{\theta}_i$. We assume they are characterized by known hyperparameters $\tilde{\beta}, \tilde{\alpha}$. Then, we can first perform the statistical inference on the surrogate parameters $\tilde{\phi}, \tilde{\theta}$. In details,

$$
\begin{aligned}
& p(r,g|c,\tilde{\beta},\tilde{\ell}) \\
= & \prod_{t=1}^{T}\int_{\tilde{\phi}_t} p(\tilde{\phi}_t|\tilde{\beta})\prod_{\tau=1}^{N_t} p(r_{i_\tau j_\tau}, g_{i_\tau j_\tau}|\tilde{\phi}_t,\tilde{\ell},c_{i_\tau j_\tau}=t)d\tilde{\phi}_t \\
= & \prod_{t=1}^{T}\int_{\phi_t} \frac{\Gamma(\sum\limits_{e=1}^{G}\tilde{\beta}_e)}{\prod\limits_{e=1}^{G}\Gamma(\tilde{\beta}_e)}\left(\prod_{e=1}^{G}\tilde{\phi}_{t,e}^{\tilde{\beta}_e-1}\right)\left(\prod_{\tau=1}^{N_t}\prod_{e=1}^{G}\tilde{\phi}_{t,e}^{\mathbb{1}\{g_{i_\tau,j_\tau}=e\}}\right)d\tilde{\phi}_t \\
= & \prod_{t=1}^{T}\frac{\Gamma(\sum\limits_{e=1}^{G}\tilde{\beta}_e)}{\prod\limits_{e=1}^{G}\Gamma(\tilde{\beta}_e)}\frac{\prod\limits_{e=1}^{G}\Gamma(\tilde{\beta}_e+\sum\limits_{\tau=1}^{N_t}\mathbb{1}\{g_{i_\tau,j_\tau}=e\})}{\Gamma(\sum\limits_{e=1}^{G}\tilde{\beta}_e+\sum\limits_{e=1}^{G}\sum\limits_{\tau=1}^{N_t}\mathbb{1}\{g_{i_\tau,j_\tau}=e\})},
\end{aligned} \tag{4}
$$

similarly, we have

$$
p(c|\tilde{\alpha},\eta)=\prod_{i=1}^{M}\frac{\Gamma(\sum\limits_{t=1}^{T}\tilde{\alpha}_t)}{\prod\limits_{t=1}^{T}\Gamma(\tilde{\alpha}_t)}\frac{\prod\limits_{t=1}^{T}\Gamma(\tilde{\alpha}_t+\sum\limits_{j=1}^{N_i}\mathbb{1}\{c_{i,j}=t\})}{\Gamma(\sum\limits_{t=1}^{T}\tilde{\alpha}_t+\sum\limits_{t=1}^{T}\sum\limits_{j=1}^{N_i}\mathbb{1}\{c_{i,j}=t\})}. \tag{5}
$$

We employed a Gibbs sampler to draw samples from the posterior distribution on the cell type assignments of all the reads from all samples. Therefore, we need the conditional distribution $p(c_a|c_{-a},r,g)$, where $r$ denotes the reads of all samples and $c_{-a}$ denotes the cell type assignments without assigning the $a^{th}$ read. In particular, assume the total number of reads is $n$, then $c=(c_1,\cdots,c_n)$ denotes the cell type assignment for all reads $r=(r_1,\cdots,r_n)$ from all the samples. To be clear, we use $gid(r_a), sid(r_a)$ as references to the gene id and sample id of read $r_a$ respectively, and $cid(r_a)$ refers to the corresponding cell type assignment used in the calculation when needed. Apply eqn.(4) and eq.(5) and the fact $\Gamma(x+1)=x\Gamma(x)$, we have

$$
\begin{aligned}
& p(c_a|c_1,\cdots,c_{a-1},c_{a+1},\cdots,c_n,r,g) \\
= & \frac{p(c_1,\cdots,c_n,r,g)}{p(c_1,\cdots,c_{a-1},c_{a+1},c_n,r,g)}
\end{aligned}
$$

$$\propto \frac{\beta_{gid(r_a)} + \sum_{\tau=1}^{N_{c_a}} \mathbb{1}_{-1}\{g_{i_\tau,j_\tau} = gid(r_a)\}}{\sum_{e=1}^{G} \beta_e + \sum_{e=1}^{G}\sum_{\tau=1}^{N_{c_a}} \mathbb{1}_{-1}\{g_{i_\tau,j_\tau} = e\}} \frac{\alpha_{c_a} + \sum_{j=1}^{N_{sid(r_a)}} \mathbb{1}_{-1}\{c_{i,j} = c_a\}}{\sum_{t=1}^{T} \alpha_t + \sum_{t=1}^{T}\sum_{j=1}^{N_{sid(r_a)}} \mathbb{1}_{-1}\{c_{i,j} = t\}}$$

where $\sum_{\tau=1}^{N_{c_a}} \mathbb{1}_{-1}\{g_{i_\tau,j_\tau} = gid(r_a)\}$ denotes the counts of all the reads with gene id $gid(r_a)$ that assigned to cell type $c_a$ without the single count of current assignment of the read $r_a$ itself. Finally, we can estimate the parameters based on the posterior predictive distribution for new data as follows,

$$\begin{aligned}
\tilde{\phi}_{t,e} &= \frac{\beta_e + \sum_{\tau=1}^{N_t} \mathbb{1}\{g_{i_\tau,j_\tau} = e\}}{\sum_{k=1}^{G} \beta_k + \sum_{k=1}^{G}\sum_{\tau=1}^{N_t} \mathbb{1}\{g_{i_\tau,j_\tau} = k\}}, \\
\tilde{\theta}_{i,t} &= \frac{\alpha_t + \sum_{j=1}^{N_i} \mathbb{1}\{c_{i,j} = t\}}{\sum_{k=1}^{T} \alpha_k + \sum_{k=1}^{T}\sum_{j=1}^{N_i} \mathbb{1}\{c_{i,j} = k\}},
\end{aligned}$$

where $t = 1, \cdots, T$ denotes cell type, $e = 1, \cdots, G$ denotes gene id, and $i = 1, \cdots, M$ denote sample id, and $\sum_{\tau=1}^{N_t} \mathbb{1}\{gid(r_{i_\tau,j_\tau}) = e\}$ is the count of reads from gene $e$ that are assigned to cell type $t$, and $\sum_{j=1}^{N_i} \mathbb{1}\{c_{i,j} = t\}$ is the count of reads from sample $i$ that are assigned to cell type $t$. Then we can recover the desired parameter $\phi, \theta$

$$\begin{aligned}
\phi_{t,e} &= \frac{\tilde{\phi}_{t,e}/\tilde{\ell}_e}{\sum_{k=1}^{G} \tilde{\phi}_{t,k}/\tilde{\ell}_k}, \\
\theta_{i,t} &= \frac{\tilde{\theta}_{i,t}/\eta_t}{\sum_{k=1}^{T} \tilde{\theta}_{i,k}/\eta_k}.
\end{aligned}$$

To infer the CDSeq-identified cell types, reference gene expression profiles of pure cell lines (raw read count preferred for RPKM normalization) is required. One can use correlation as a metric to associate the cell type with the cell types in the reference profile.

**Determine the number of cell types using the data** Our method allows the number of cell types, which in some cases may be unknown, to be inferred from the data. Using our statistical model, assuming the hyperparameters $\alpha, \beta$ and the reads mapping information are known. Let $\mathcal{C}$ denote the space of all possible cell type assignments for all the reads, then summing over $c \in \mathcal{C}$, the joint probability distribution of the cell type-specific gene expression profiles, the sample-specific proportions of the cell types, and the mapped reads is given in (6).

$$
\begin{aligned}
& p(r, g, \theta, \phi | \alpha, \beta, \eta, \tilde{\ell}) \\
= & \sum_{c \in \mathcal{C}} p(r, g, c, \theta, \phi | \alpha, \beta, \eta, \tilde{\ell}) \\
= & \sum_{c \in \mathcal{C}} \prod_{t=1}^{T} p(\phi_t | \beta) \prod_{i=1}^{M} \left( p(\theta_i | \alpha) \prod_{j=1}^{N_i} p(c_{i,j} | \theta_i, \eta) p(r_{i,j}, g_{i,j} | \phi, \tilde{\ell}, c_{i,j}) \right) \\
= & \prod_{t=1}^{T} p(\phi_t | \beta) \prod_{i=1}^{M} \left( p(\theta_i | \alpha) \prod_{j=1}^{N_i} \sum_{c \in \mathcal{C}} p(c_{i,j} | \theta_i, \eta) p(r_{i,j}, g_{i,j} | \phi, \tilde{\ell}, c_{i,j}) \right) \\
= & \prod_{t=1}^{T} p(\phi_t | \beta) \prod_{i=1}^{M} \left( p(\theta_i | \alpha) \prod_{j=1}^{N_i} \sum_{c_{i,j}=1}^{T} p(c_{i,j} | \theta_i, \eta) p(r_{i,j}, g_{i,j} | \phi, \tilde{\ell}, c_{i,j}) \right) \\
= & \prod_{t=1}^{T} p(\phi_t | \beta) \prod_{i=1}^{M} \left( p(\theta_i | \alpha) \prod_{j=1}^{N_i} \sum_{c_{i,j}=1}^{T} \frac{\theta_{i,c_{i,j}} \eta_{c_{i,j}}}{\sum_{t=1}^{T} \theta_{i,t} \eta_t} \frac{\phi_{c_{i,j}, g_{i,j}} \tilde{\ell}_{g_{i,j}}}{\sum_{k=1}^{G} \phi_{t,k} \tilde{\ell}_k} \right) \\
= & \prod_{t=1}^{T} p(\phi_t | \beta) \prod_{i=1}^{M} \left( p(\theta_i | \alpha) \prod_{e=1}^{G} \left( \sum_{t=1}^{T} \frac{\theta_{i,t} \eta_t}{\sum_{\tau=1}^{T} \theta_{i,\tau} \eta_\tau} \frac{\phi_{t,e} \tilde{\ell}_e}{\sum_{k=1}^{G} \phi_{t,k} \tilde{\ell}_k} \right)^{n_{i,e}} \right) \\
= & \prod_{t=1}^{T} \frac{\Gamma(\sum_{e=1}^{G} \beta_e)}{\prod_{e=1}^{G} \Gamma(\beta_e)} \left( \prod_{e=1}^{G} \phi_{t,e}^{\beta_e - 1} \right) \prod_{i=1}^{M} \left( \frac{\Gamma(\sum_{t=1}^{T} \alpha_t)}{\prod_{t=1}^{T} \Gamma(\alpha_t)} \left( \prod_{t=1}^{T} \theta_{i,t}^{\alpha_t - 1} \right) \prod_{e=1}^{G} \left( \sum_{t=1}^{T} \frac{\theta_{i,t} \eta_t}{\sum_{\tau=1}^{T} \theta_{i,\tau} \eta_\tau} \frac{\phi_{t,e} \tilde{\ell}_e}{\sum_{k=1}^{G} \phi_{t,k} \tilde{\ell}_k} \right)^{n_{i,e}} \right)
\end{aligned}
$$

$$\propto \prod_{t=1}^{T}\left(\prod_{e=1}^{G}\phi_{t,e}^{\beta_e-1}\right)\prod_{i=1}^{M}\left(\left(\prod_{t=1}^{T}\theta_{i,t}^{\alpha_t-1}\right)\prod_{e=1}^{G}\left(\sum_{t=1}^{T}\frac{\theta_{i,t}\eta_t}{\sum_{\tau=1}^{T}\theta_{i,\tau}\eta_\tau}\frac{\phi_{t,e}\tilde{\ell}_e}{\sum_{k=1}^{G}\phi_{t,k}\tilde{\ell}_k}\right)^{n_{i,e}}\right), \tag{6}$$

where $n_{i,e}$ denotes the count of the reads mapped to gene $e$ in sample $i$. Then taking logarithmic of the posterior in (6), we have

$$\begin{aligned}
&\log p(r,\theta,\phi|\alpha,\beta,\eta,\ell,g)\\
=\;& \sum_{t=1}^{T}\sum_{e=1}^{G}(\beta_e-1)\log(\phi_{t,e})+\sum_{i=1}^{M}\left(\sum_{t=1}^{T}(\alpha_t-1)\log\theta_{i,t}+\sum_{e=1}^{G}n_{i,e}\log\left(\sum_{t=1}^{T}\frac{\theta_{i,t}\eta_t}{\sum_{\tau=1}^{T}\theta_{i\tau}\eta_\tau}\frac{\phi_{t,e}\tilde{\ell}_e}{\sum_{k=1}^{G}\phi_{t,k}\tilde{\ell}_k}\right)\right)\\
=\;& \underbrace{\sum_{t=1}^{T}\sum_{e=1}^{G}(\beta_e-1)\log(\phi_{t,e})+\sum_{i=1}^{M}\sum_{t=1}^{T}(\alpha_t-1)\log\theta_{i,t}}_{\text{regulation terms imposed by prior distributions}}+\underbrace{\sum_{i=1}^{M}\sum_{e=1}^{G}n_{i,e}\log\left(\sum_{t=1}^{T}\frac{\theta_{i,t}\eta_t}{\sum_{k=1}^{T}\theta_{i,k}\eta_k}\frac{\phi_{t,e}\tilde{\ell}_e}{\sum_{k=1}^{G}\phi_{t,k}\tilde{\ell}_k}\right)}_{\text{likelihood of the data}}.
\end{aligned} \tag{7}$$

The log posterior of RNA-seq data given in (7) can be simplified as follows

$$\begin{aligned}
&h(\tilde{\theta},\tilde{\phi},r)\\
=\;& \sum_{t=1}^{T}\sum_{e=1}^{G}(\tilde{\beta}_e-1)\log(\tilde{\phi}_{t,e})+\sum_{i=1}^{M}\sum_{t=1}^{T}(\tilde{\alpha}_t-1)\log\tilde{\theta}_{i,t}+\sum_{i=1}^{M}\sum_{e=1}^{G}n_{i,e}\log\left(\sum_{t=1}^{T}\tilde{\theta}_{i,t}\tilde{\phi}_{t,e}\right),
\end{aligned} \tag{8}$$

Given a set of candidate values for the number of cell types present in the mixture samples, CDSeq will choose the one that maximizes eqn (8).
